# Loss of Ferroptosis-suppressors Accelerates Breast Cancer Recurrence by Promoting Dormancy Exit

**DOI:** 10.64898/2026.10.22.761187

**Authors:** Emily K. Shea, Matthew Lawrence-Paul, Amulya Sreekumar, Brian Benz, Christopher Sterner, Jianping Wang, Yan Chen, Tien-chi Pan, George Belka, Lewis A. Chodosh

## Abstract

Breast cancer mortality is principally due to recurrent metastatic disease that may commonly emerge from a state of dormant minimal residual disease following definitive treatment of a primary tumor. Our genomic analysis of paired primary and metastatic breast cancers arising in the same patients resulted in the identification of the ferroptosis pathway as being preferentially mutated in metastases. An in vivo CRISPR-Cas9 screen of known ferroptosis regulators in mice identified guides targeting *PNPLA2*, *GOT1*, and *GPX4* as being selected for in late dormancy and recurrence, which was surprising given the ferroptosis-suppressing functions ascribed to these proteins. However, recurrence-free survival assays in mice confirmed that deletion of *PNPLA2*, *GOT1*, or *GPX4* accelerated tumor recurrence in mice and provided further evidence that this acceleration was attributable to an increased probability that dormant tumor cells deleted for these genes would reenter the cell cycle. Using a model of therapy-associated breast cancer dormancy, we also found that dormant cancer cells were markedly resistant to ferroptosis-inducing stimuli, including erastin and RSL3. Moreover, dormant tumor cells lacking *PNPLA2*, *GOT1*, or *GPX4* displayed enhanced ferroptosis resistance compared to wildtype cells. Probing this ferroptosis-resistant state revealed that the ratio of reduced to oxidized glutathione was elevated in dormant cells. Moreover, deletion of *PNPLA2*, *GOT1*, or *GPX4* further increased this ratio in dormant cells, suggesting that an increased ratio of reduced to oxidized glutathione may contribute to ferroptosis resistance in dormant tumor cells, especially those lacking these genes, potentially by rendering them more capable of handling lipid oxidative stress. Consistent with our findings in mice, analysis of human breast cancer patient data revealed that patients with copy number loss of *PNPLA2*, *GOT1*, or *GPX4* exhibited worse recurrence-free and overall survival. Together, our study implicates three genes in the process of recurrence from dormant tumor cells and identifies a unique metabolic feature of the dormant tumor cell state. The surprising resistance to ferroptosis inducers exhibited by dormant tumor cells, as well as counterintuitive effects of deleting ferroptosis-suppressing genes on tumor recurrence following dormancy, raise questions regarding the utility of targeting ferroptosis in cancer that merit further study.

## Introduction

Breast cancer is the most prevalent cancer worldwide and leading cause of cancer death among women, with mortality principally being attributable to recurrent metastatic disease (1–3). A unique feature of breast cancer is the relatively constant risk of recurrence that persists for decades following primary tumor therapy, particularly for estrogen receptor-positive (ER+) disease (4). The extended periods of clinical latency that often occur between definitive treatment of primary tumors and their eventual recurrence suggest that residual tumor cells (RTCs) surviving adjuvant therapy may pass through a dormant state, either due to reversible exit from the cell cycle, referred to as cellular dormancy, or balanced rates of tumor cell proliferation and death, referred to as tumor mass dormancy (5–8).

Our laboratory has previously generated and extensively validated genetically engineered mouse models of breast cancer dormancy and recurrence. Employing a mammary-specific TetOn system, the presence of doxycycline triggers expression of a driver oncogene relevant to human breast cancer (e.g., Her2, Wnt1, Akt1, Myc) specifically in the mammary epithelium, ultimately resulting in the development of an invasive primary tumor (9–13).

Following primary tumor formation, withdrawal of doxycycline treatment results in the downregulation of the driver oncogene and regression of primary tumors to a non-palpable state characterized by dormant minimal residual disease. These populations of dormant RTCs spontaneously give rise to recurrent tumors over the course of weeks to months. Further, primary tumor cells from the inducible Her2 model have been isolated and cultured to enable these cells to be genetically modified and then injected orthotopically in vivo or studied in vitro (14–16). These model systems allow the mechanisms underlying dormancy and recurrence to be systematically analyzed.

Ferroptosis is a form of non-apoptotic cell death (17) whose three key requirements are iron, oxidizable lipids (e.g., polyunsaturated fatty acids: PUFAs), and a relative lack of oxidative damage repair capacity (18). Beyond these essential requirements, ferroptosis is impacted by a number of other cellular pathways, including iron and lipid metabolism, as well as redox homeostasis (19,20). Given the well-described ability of cancer cells to evade apoptosis, ferroptosis has become an attractive field for investigating potential cancer cell vulnerabilities that could be leveraged therapeutically. Indeed, several studies have shown promise in targeting ferroptosis as a therapy for cancer (18,21,22).

To date, the role of ferroptosis in cancer has principally been studied in the context of primary tumor growth. Intriguingly, two recent studies have implicated agents that cause cell cycle arrest and microenvironment-induced dormancy as potential causes of ferroptosis resistance in cancer cells (23,24). Herein, we explore whether ferroptosis sensitivity is altered during dormancy, and whether genes known to modulate ferroptosis sensitivity impact cellular dormancy and tumor recurrence.

## Results

### An in vivo CRISPR screen counterintuitively identifies selection for the deletion of ferroptosis-suppressing genes during tumor recurrence

In a large cohort (n=130) of paired primary and metastatic breast cancers, we found that the ferroptosis pathway was preferentially mutated in metastatic tumors compared to the paired primary tumors from which they arose (13% of metastases, 2% of primary tumors, p = 0.0034) (Fig. 1a). The nature of this analysis, however, did not permit us to infer whether metastasis-specific mutations resulted in ferroptosis activation or inhibition. Therefore, to explore the hypothesis that ferroptosis dysregulation may contribute to cancer dormancy and/or recurrence, we conducted a targeted CRISPR screen.

**Figure 1:**
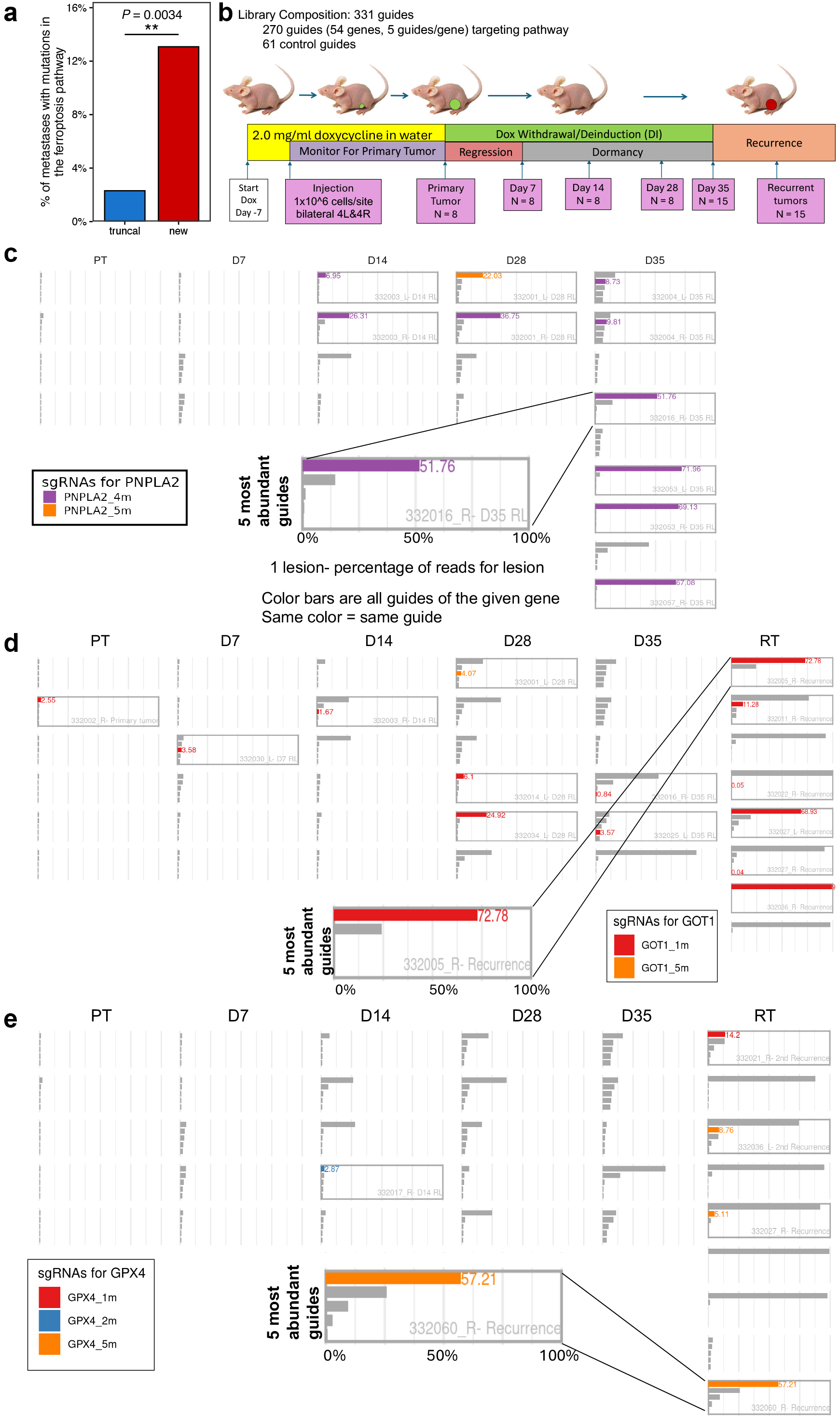
An in vivo CRISPR screen counterintuitively identifies selection for the deletion ferroptosis-suppressing genes during tumor recurrence a. Percentage of tumors harboring a mutation in a ferroptosis-related gene comparing metastases (new) vs. primary tumors (truncal) based on WES in 130 paired tumors. P-value from one-sided Wilcoxon signed-rank test. b. Schematic of CRISPR Library Design: 331 guides consisting of 5 guides/gene for 54 genes of interest and 61 control guides were transduced into iHer2 cells and injected orthotopically into nude mice. c,d,e. Barplots showing the top 5 guides per lesion, with highlighted bars representing guides targeting *PNPLA2*, *GOT1*, and *GPX4*, respectively.

We designed a library of CRISPR guides targeting genes reported to play a functional role in ferroptosis. This library consisted of 331 guides, including 5 guides targeting each of 54 genes, along with control guides, in a lentiviral vector backbone containing a GFP marker (25) (Table 1). Inducible Her2 primary tumors cells previously isolated and cultured from MMTV-rtTA;TetO-Her2/neu (MTB/TAN) mice (10,14) (hereafter referred to as iHer2 cells) stably expressing Cas9 were then transduced with this library at a multiplicity of infection (MOI) of 0.3 to ensure each cell integrated no more than one guide. Following cell sorting for GFP expression, cells were expanded and injected orthotopically into the mammary fat pads of nu/nu mice maintained on doxycycline (Fig. 1b). Following primary tumor growth, doxycycline was withdrawn to induce tumor regression to a dormant state and then monitored for spontaneous tumor recurrence, as previously described (14,16,26). Primary tumors, dormant residual lesions at days 7, 14, 28, and 35 post-doxycycline withdrawal, and recurrent tumors were collected and sequenced to determine guide representation.

**Table 1:**
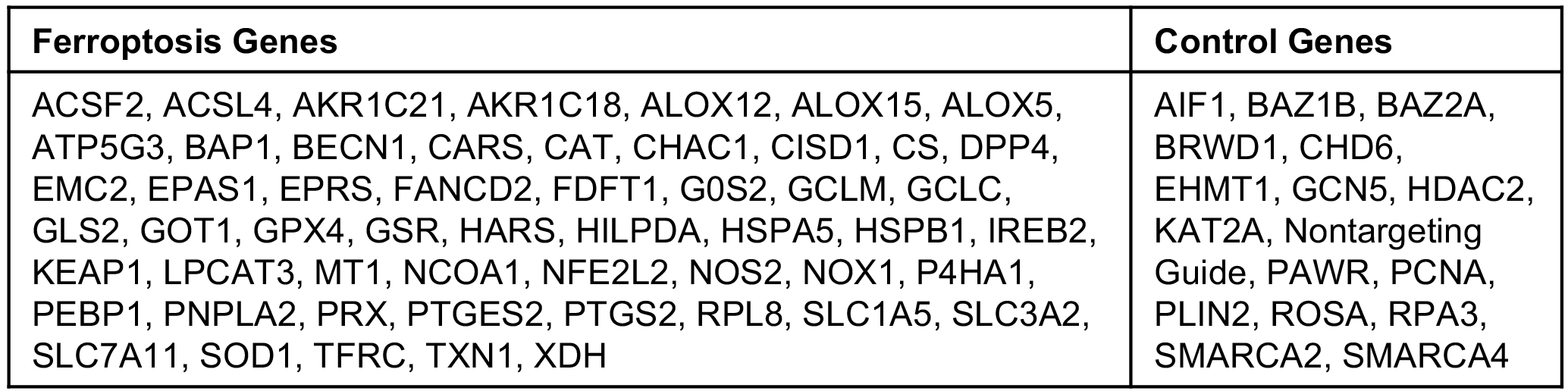
The 54 genes included in the CRISOR-cas9 screen.

| **Ferroptosis Genes** | **Control Genes** |
| --- | --- |
| ACSF2, ACSL4, AKR1C21, AKR1C18, ALOX12, ALOX15, ALOX5, ATP5G3, BAP1, BECN1, CARS, CAT, CHAC1, CISD1, CS, DPP4, EMC2, EPAS1, EPRS, FANCD2, FDFT1, G0S2, GCLM, GCLC, GLS2, GOT1, GPX4, GSR, HARS, HILPDA, HSPA5, HSPB1, IREB2, KEAP1, LPCAT3, MT1, NCOA1, NFE2L2, NOS2, NOX1, P4HA1, PEBP1, PNPLA2, PRX, PTGES2, PTGS2, RPL8, SLC1A5, SLC3A2, SLC7A11, SOD1, TFRC, TXN1, XDH | AIF1, BAZ1B, BAZ2A, BRWD1, CHD6, EHMT1, GCN5, HDAC2, KAT2A, Nontargeting Guide, PAWR, PCNA, PLIN2, ROSA, RPA3, SMARCA2, SMARCA4 |

We anticipated that deletion of a gene that suppresses recurrence would give rise to a clonal outgrowth of tumor cells. Three methods were used to identify and rank clonal hits: 1) abundance, evaluated based on how many neoplastic lesions across the screen exhibited 5% or greater reads for a specific guide; 2) outliers, evaluated based on how many lesions across the screen exhibited reads for a specific guide greater than 1.5x the interquartile range; and 3) significance, evaluated based on how many lesions across the screen exhibited a greater than 2-fold increase in reads compared with primary tumors, along with an adjusted p-value < 0.1 (Supp. Fig. 1a-c).

To validate these analytical methods clonal hits were also compared to a negative control library screen, where every guide in the library was non-targeting. In quantifying results from the negative control library, the mean number of lesions with high abundance was 0.4 (standard deviation = 0.6), the mean number of outlier lesions was 1.7 (standard deviation = 1.9), and the mean number of significant lesions was 5.9 (standard deviation = 4.4).

*PNPLA2* was identified as the top hit from the screen as measured by guide abundance, with sg*PNPLA2* guides comprising >5% of reads in 10 dormant residual lesions and early recurrences out of 63 total lesions analyzed across all timepoints. Indeed, sg*PNPLA2* was the most abundant guide in 9 of these 10 lesions (Fig. 1c, Supp. Fig. 1a). sg*PNPLA2* was also ranked among the top three hits when ranked either by number of outliers (16 lesions) or by significance (24 lesions).

*PNPLA2* encodes the enzyme adipose triglyceride lipase (ATGL), which catalyzes the initial step of triglyceride hydrolysis (27,28). Given its role in releasing fatty acids, ATGL has been proposed to have both ferroptosis promoting and ferroptosis inhibiting roles. Specifically, the release of polyunsaturated fatty acids (PUFAs) promotes lipid peroxidation and ferroptosis, whereas release of monounsaturated fatty acids (MUFAs) suppresses ferroptosis (29).

Ferroptosis suppression in this context is thought to be mediated by the displacement of PUFAs in the plasma membrane by MUFAs, thereby resulting in a reduction in oxidized lipids (30) and/or activation of PPAR-α signaling, which upregulates the cell’s oxidative capacity (31).

The second-ranked gene by abundance in our CRISPR screen was *GOT1.* sg*GOT1* was selected for in four out of 15 recurrent tumors, within which it represented the dominant clone (Fig. 1d, Supp. Fig. 1b). sg*GOT1* was also the top-ranked guide when assessed by number of outliers (20 lesions), and the sixth-ranked gene when assessed by significance (22 lesions).

*GOT1* encodes glutamic-oxaloacetic transaminase 1, which catalyzes the reversible conversion of glutamate and oxaloacetate to aspartate and alpha-ketoglutarate (32). GOT1 plays a role in multiple cellular metabolic pathways including, but not limited to, NADPH synthesis, nucleotide synthesis, and glycolysis regulation (33). In particular, GOT1 has been reported to protect against ferroptosis by suppressing the breakdown of ferritin, thereby reducing the labile iron pool and preventing stimulation of ferroptotic death (34). Further, GOT1 inhibition has also been shown to reduce NADPH and GSH levels, resulting in increased ferroptosis (34).

*GPX4* was ranked sixth in our CRISPR screen when evaluated by abundance and its selection in four recurrent tumors was particularly surprising given its key role in preventing ferroptotic cell death (Fig. 1e, Supp. Fig. 1c) (35). By both outlier analysis (7 lesions) and significance (17 lesions), sg*GPX4* was ranked 11^th^, which is significant when compared to findings from the negative control library. Results for *PNPLA2*, *GOT1*, *GPX4* using each of the three metrics employed are summarized in Table 2.

**Table 2:**
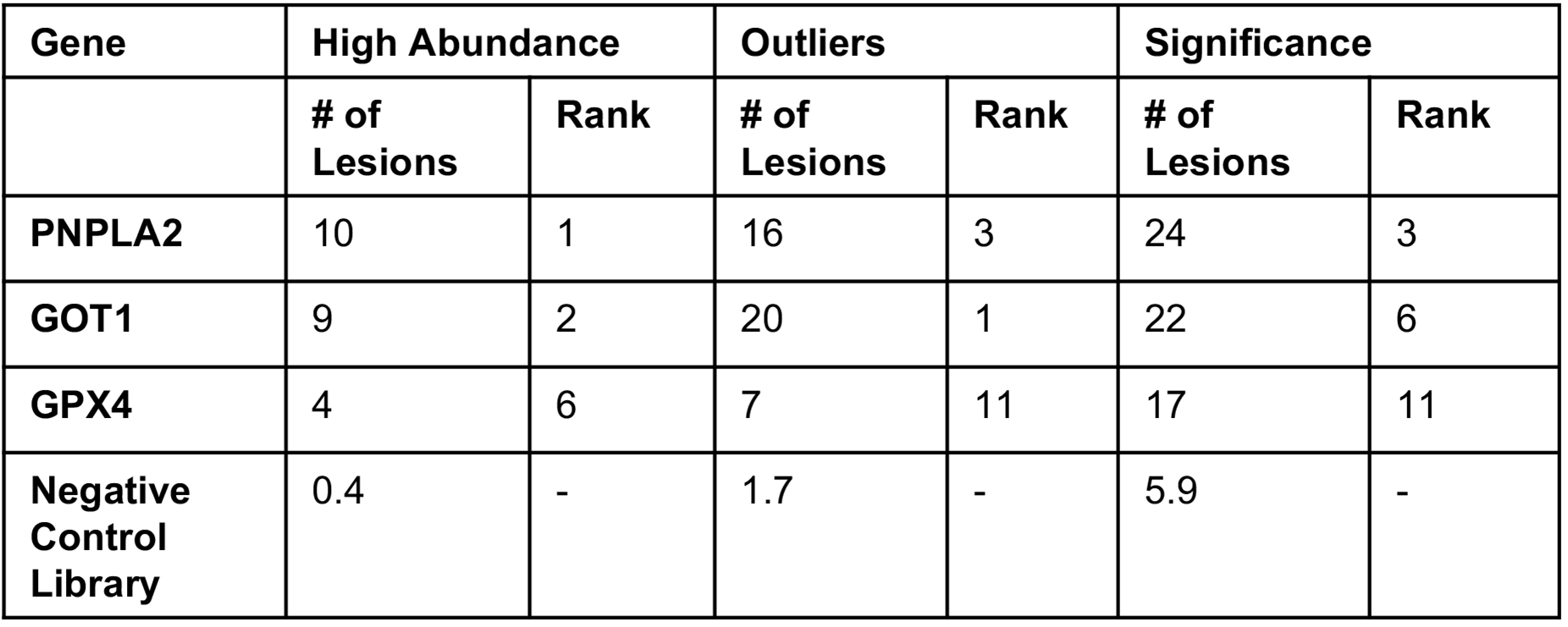
Summarized high abundance outliers, and significance scores and ranks for PNPLA2, GOT1, and GPX4 from the CRISP-cas9 screen, as compared to the negative control library.

*GPX4* encodes glutathione peroxidase 4, one of eight glutathione peroxidase enzymes (36). Notably, it is the only glutathione peroxidase that can reduce lipid peroxides in membranes. Using reduced glutathione, GPX4 catalyzes the reduction of lipid peroxides to non-toxic lipid alcohols, which constitutes a canonical ferroptosis-suppression mechanism (37).

As PNPLA2, GOT1 and GPX4 each have known roles in preventing ferroptotic cell death, we anticipated that guides targeting these genes would increase susceptibility to ferroptotic cell death and, consequently, would be selected against during tumor dormancy and recurrence. Counterintuitively, we found that each of these guides was enriched in late dormancy and recurrence.

### Deletion of *PNPLA2*, *GOT1*, and *GPX4* each accelerate recurrence

To confirm that each of the guides was effective against its intended target, TIDE analysis (38,39) was performed on cells transduced with the guides at an MOI of 3. Both *PNPLA2* guides, *PNPLA2g4* and *PNPLA2g5*, exhibited high indel percentage and knockout scores (*PNPLA2g4* indel % = 87%, knockout score = 87; *PNPLA2g5* indel % = 84, knockout score = 76) (Supp. Fig. 2a, b). TIDE analysis similarly revealed that the most highly enriched guides for GOT1 and GPX4 from the screen each targeted its intended gene of interest (*GOT1g1*: indel % = 61, knockout score = 54; *GPX4g5*: indel % = 80, knockout score = 67) (Supp. Fig. 2c, e).

**Figure 2:**
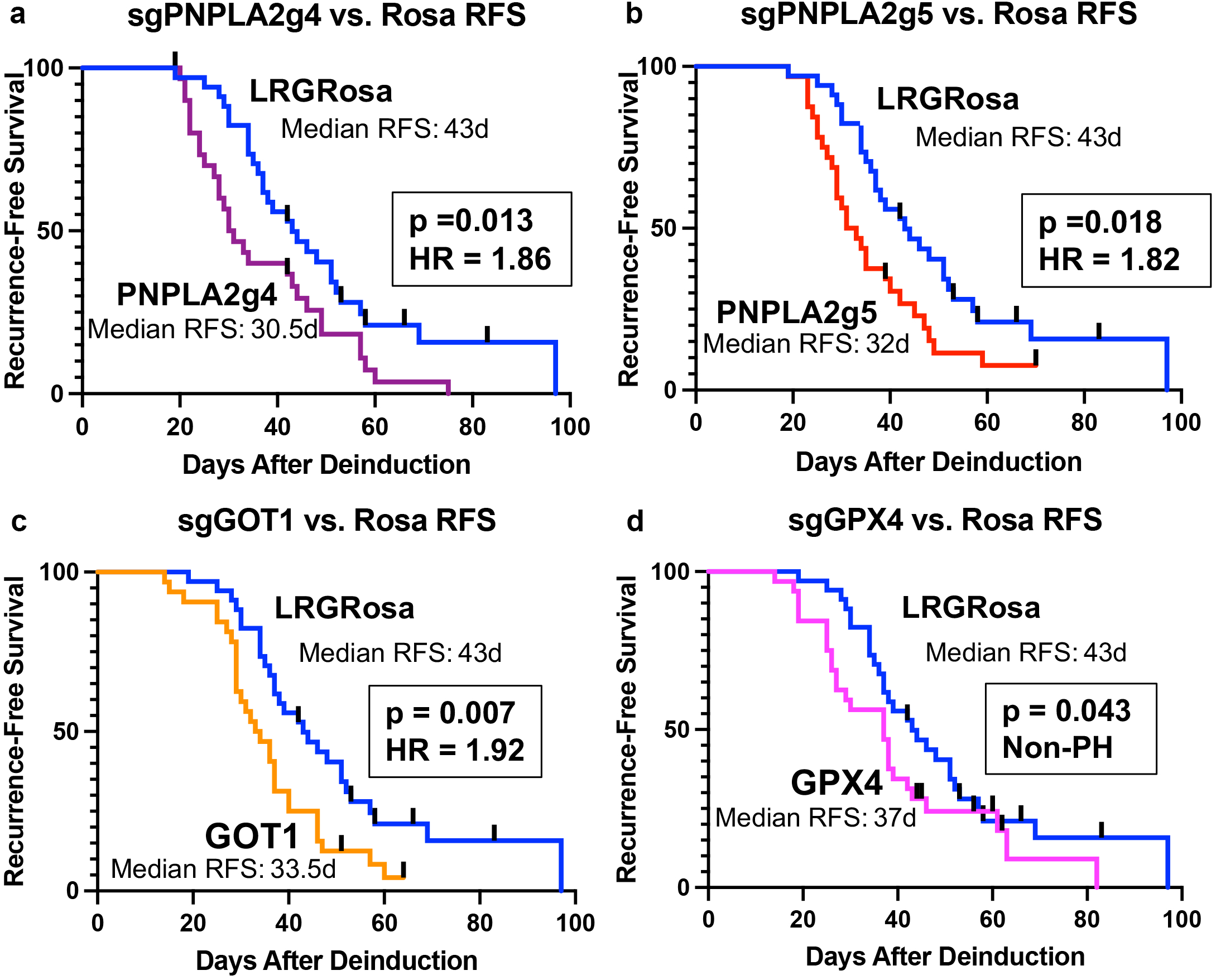
Deletion of ferroptotic regulators accelerates recurrence Kaplan-Meier curves for: a. *PNPLA2g4* b. *PNPLA2g5* c. *GOT1g1* d. *GPX4g5* showing recurrence-free survival compared to sg*Rosa* control. Hazard ratios and p-values calculated from logrank test, except in the case of *GPX4g5*, where curves crossed and p-value is calculated by a weighted log rank test for early difference.

To determine directly whether deletion of *PNPLA2*, *GOT1*, or *GPX4* promoted tumor recurrence, 16 nu/nu mice per arm were injected orthotopically with Cas9-expressing iHer2 cells transduced with a guide targeting *GOT1*, *GPX4*, *PNPLA2*, or the *Rosa* locus. After allowing primary tumors to form, doxycycline treatment was discontinued to downregulate Her2 expression and induce tumor regression to a non-palpable state. Mice were monitored three times weekly for spontaneous tumor recurrence.

Consistent with our findings in the CRISPR screen, each of two guides targeting *PNPLA2* significantly accelerated recurrence compared to sg*Rosa* control (logrank HR = 1.86 and 1.82, p = 0.013 and 0.018, respectively) (Fig. 2a, b). Compared with the median time to recurrence in the control arm of 43 days, each of the two *PNPLA2* guides tested significantly decreased the median time to recurrence to 30.5 and 32 days, respectively.

Further confirming results of the CRISPR screen, deletion of *GOT1* also significantly accelerated recurrence (HR = 1.92, p = 0.007), while reducing median time to recurrence to 33.5 days (Fig. 2c). *GPX4* deletion exerted a more modest effect on recurrence, reducing median time to recurrence to 37 days (p = 0.043, non-proportional hazards) (Fig. 2d).

An additional in vivo competition experiment performed with mixtures of 20% sg*PNPLA2*, sg*GOT1*, or sg*GPX4* cells co-injected with 80% sg*Rosa* cells confirmed the acceleration of recurrence resulting from deletion of each of these genes (Supp. Fig. 3). In addition to the original guides used in the recurrence-free survival analysis, *GOT1g5* was used as an additional guide that was selected for in the screen, whereas *GPX4g8* was newly designed. Both were confirmed to target their intended gene by TIDE analysis (Supp. Fig. 2d, f).

**Figure 3:**
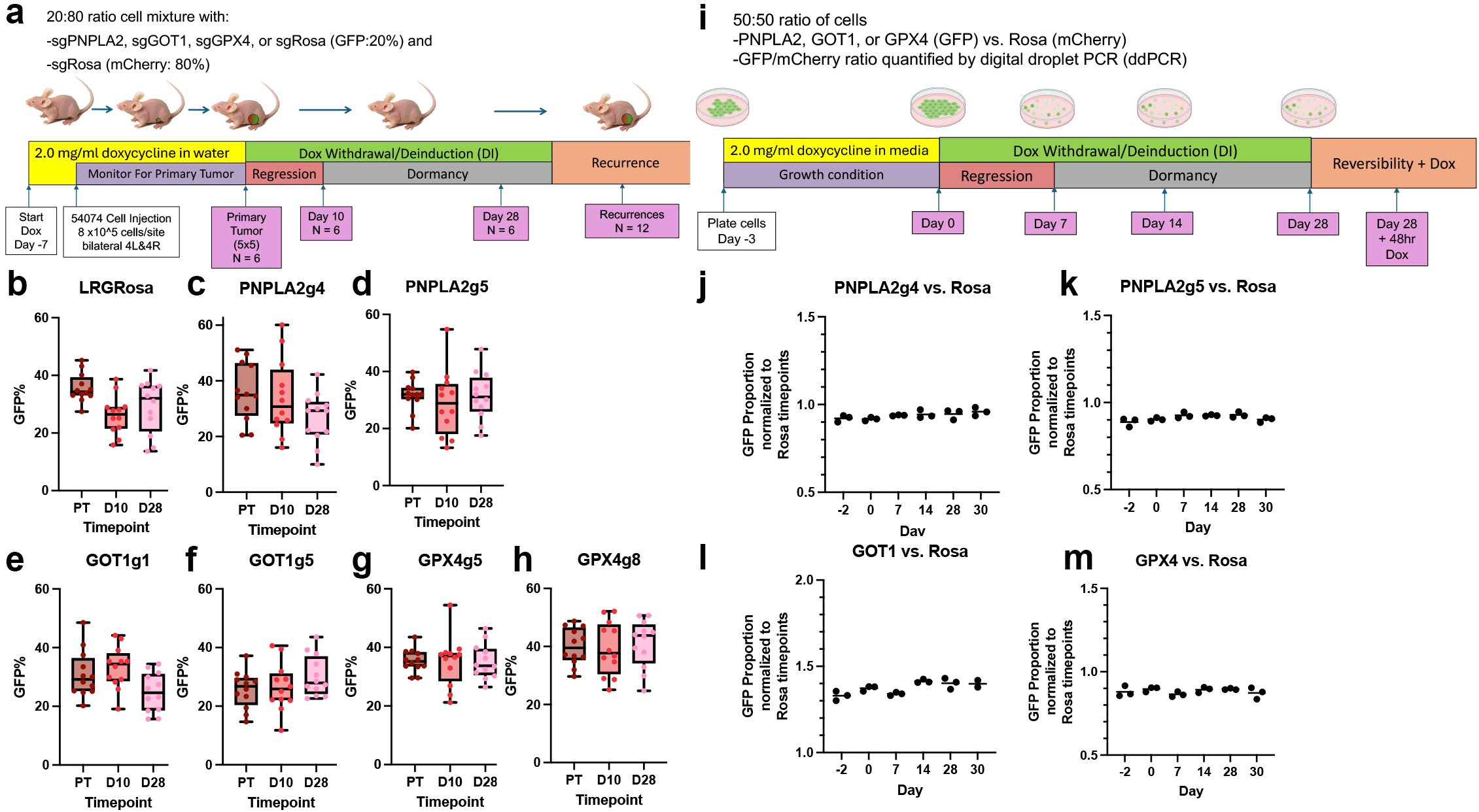
Deletion of ferroptotic regulators does not alter cellular fitness during dormancy a. Schematic of in vivo competition assay, showing timepoints collected, cell conditions and state at each time point, and cell ratio. b-h. GFP% of cell competition at timepoint listed. GFP guide was b. sg*Rosa*, c. *PNPLA2g4*, d. *PNPLA2g5*, e. *GOT1g1*, f. *GOT1g5*, g. *GPX4g5*, h. *GPX4g8*. i. Schematic of in vitro competition assay, showing timepoints collected, cell conditions and state at each time point, and cell ratio. j-m. GFP proportion of cell competition at indicated timepoints. Proportion has been normalized to proportion of GFP at the timepoint listed in the Rosa-GFP vs. Rosa-mCherry control arm. GFP guide was j. *PNPLA2g4*, k. *PNPLA2g5*, l. *GOT1g1*, m. *GPX4g5*.

*PNPLA2g4* and *PNPLA2g5* reduced median RFS to 35 and 42 days, respectively, as compared to 50 days in the Rosa control group (Supp. Fig. 3a). Guide 4 exhibited a statistically significant acceleration (HR = 3.3, p < 0.0001), whereas guide 5 trended towards significance (HR = 1.72, p = 0.11). For *GOT1*, both guide 1 (HR = 3.26, p < 0.0001) and guide 5 (HR = 3.27, p < 0.0001) significantly accelerated recurrence by reducing median RFS to 35 days (Supp. Fig. 3b). Further, *GPX4g5* (HR = 1.94, p = 0.041) and *GPX4g8* (HR = 2.27, p = 0.010) significantly accelerated recurrence, reducing median RFS to 36 and 35 days, respectively (Supp. Fig. 3c).

Together, the acceleration in recurrence that we observed in mice for guides targeting *PNPLA2*, *GOT1*, and *GPX4* was consistent with the counterintuitive findings from our CRISPR screen. Whereas deletion of each of these ferroptosis-suppressing genes would have been anticipated to increase susceptibility to ferroptotic cell death, resulting in fewer residual tumor cells and delayed tumor recurrence, we instead observed that deletion of *PNPLA2*, *GOT1* or *GPX4* each resulted in accelerated tumor recurrence.

### Deletion of *PNPLA2*, *GOT1*, or *GPX4* does not Alter Cellular Fitness during Dormancy

A potential explanation for how deletion of *PNPLA2, GOT1,* or *GPX4* could accelerate tumor recurrence would be if deletion resulted in increased cellular fitness during dormancy, such that a greater number of surviving dormant residual tumor cells would provide a greater number of cells from which recurrent tumors could arise. If true, we predicted that we would observe increasing selection over time for cells with deletion of each of these genes during the dormancy time course.

To test this hypothesis, we orthotopically injected nu/nu mice maintained on doxycycline with mixtures of iHer2 cells where 20% of cells contained a guide of interest and 80% of cells contained a guide targeting *Rosa*. Experimental arms included two different guides for each gene of interest (*GOT1*, *GPX4*, and *PNPLA2*) in addition to a *Rosa* vs. *Rosa* control group.

Primary tumors (PT), early (D10) and late (D28) dormant residual lesions were collected following Her2 downregulation (Fig. 3a). Following DNA isolation, the ratio of cell genotypes within each lesion was determined by droplet digital PCR (ddPCR) for GFP (in the lentiviral vector expressing the guides of interest) and mCherry (in the lentiviral vector expressing the *Rosa* guide).

Across genotypes, we did not observe selection for or against guides targeting *PNPLA2*, *GOT1*, and *GPX4* either following acute Her2 downregulation (PT to D10) or as function of time during dormancy (D10 to D28) (Fig. 3c-h). Indeed, the only selection observed was in the *Rosa* vs. *Rosa* control group for the PT to D10 interval (p = 0.017, Bonferroni correction for 21 tests) (Fig. 3b). Strikingly, none of the comparisons (PT to D10, D10 to D28, or PT to D28) were significantly different within any of the experimental arms (Fig. 3c-h). These findings suggest that from primary tumor formation through D28 of dormancy, changes in cell fitness are not evident following deletion of *PNPLA2, GOT1,* or *GPX4*.

To further investigate this hypothesis, we performed an in vitro competition assay where iHer2 cells bearing a deleted gene of interest were admixed in a 1:1 ratio with iHer2 cells containing a guide targeting *Rosa*. The mixed cell population was induced to enter a dormant state by Her2 downregulation triggered by doxycycline withdrawal. Cell populations were harvested 2 days prior to doxycycline withdrawal (D-2), on the day of doxycycline withdrawal (D0), or 7 (D7), 14 (D14) or 28 (D28) days following doxycycline withdrawal (Fig. 3i).

For each of the three gene deletions evaluated, including two guides targeting *PNPLA2* and one guide each targeting *GOT1* and *GPX4*, no significant changes were observed in the ratio of cells from D-2 to D28 (Fig. 3j-m). This result suggests an absence of selection for, or against, tumor cells harboring deletions in *PNPLA2*, *GOT1* or *GPX4* following acute Her2 downregulation or during dormancy. The lack of selection for tumor cells transduced with guides targeting *PNPLA2*, *GOT1*, or *GPX4* during dormancy, both in vivo and in vitro, argues against the hypothesis that deletion of each of these genes accelerates recurrence by increasing cellular fitness during dormancy.

### Deletion of *PNPLA2*, *GOT1*, or *GPX4* does not Alter the Rate of Tumor Growth

A second potential explanation for how gene deletion could accelerate tumor recurrence would be if gene deletion resulted in an increase in the rate of cell proliferation in dormant tumor cells that had re-entered the cell cycle, thereby increasing the rate of recurrent tumor growth and decreasing the latency to clinical detection by palpation. We therefore employed two methods to estimate tumor growth rate: linear growth rate and volume doubling rate.

No statistically significant changes in recurrent tumor growth rates were observed by either method for sg*PNPLA2*, sg*GOT1*, or sg*GPX4* recurrent tumors compared to sg*Rosa* control tumors (Fig. 4a, b). This suggests that differences in time to recurrence were not due to faster tumor growth rates once dormant cells had reentered the cell cycle. In fact, sg*GOT1* recurrent tumors exhibited slightly slower recurrent tumor growth rates compared to sg*Rosa* recurrent tumors, which would not explain accelerated recurrence.

**Figure 4:**
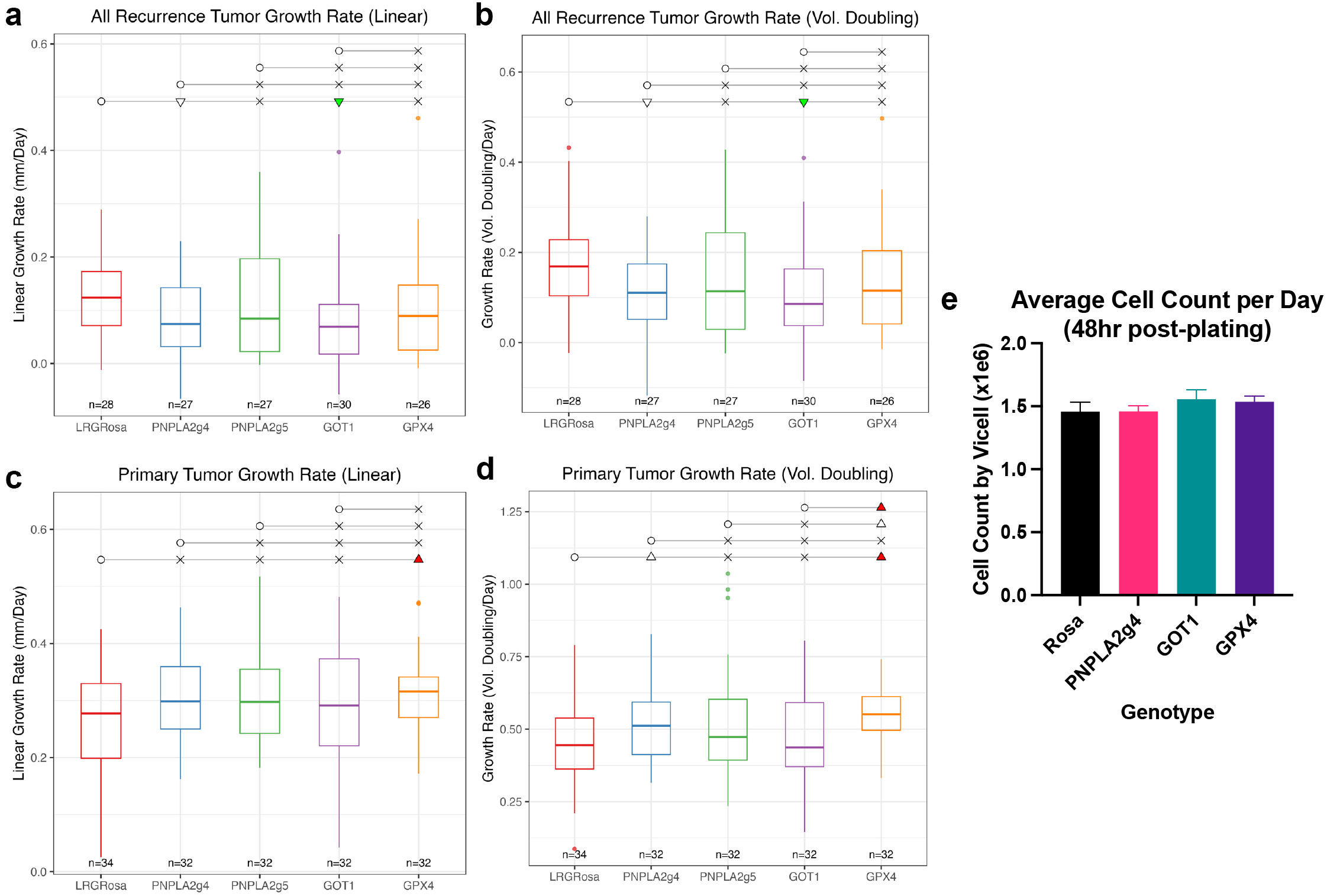
Deletion of ferroptotic regulators does not change tumor growth rate Tumor growth rates calculated from recurrence-free survival experiment in Fig 2: a. Rates of recurrent tumor growth calculated by linear growth rate from date of recurrence to date of sacrifice; b. Rates of recurrent tumor growth calculated by volume doubling rate from date of recurrence to date of sacrifice; c. Rates of primary tumor growth calculated by linear growth rate from date of injection to date of 5mm x 5mm primary tumor size; d. Rates of primary tumor growth calculated by volume doubling rate from date of injection to date of 5mm x 5mm primary tumor size. Pairwise comparisons are indicated by horizontal lines where an open circle represents the comparison timepoint, an “X” indicates no significant difference, an open triangle indicates a trending difference in the direction it points, a green triangle indicates a statistically significant decrease and a red triangle indicates a statistically significant increase. e. In vitro average cells counts per day per genotype represented as of 7 every-other day counts over the course of 14 days.

Further support for this result came from measurements of primary tumors, where growth rates were calculated from the time of injection to de-induction when primary tumors had reached a target size of 5mm x 5mm. Primary tumor growth rates were similar for sg*PNPLA2*, sg*GOT1*, and sg*Rosa* controls, and only marginally faster for sg*GPX4* tumors as estimated by each method. (Fig. 4c, d).

As a further test of the similarity in growth rates for sg*PNPLA2*, sg*GOT1*, and sg*GPX4* tumor cells compared to sg*Rosa* controls, we measured in vitro proliferation rates. Cells transduced with guides targeting *GPX4*, *GOT1*, *PNPLA2*, or *Rosa* were counted and replated every other day for two weeks. The average cell count per day across the time course was consistent across all groups, indicating that none of these gene deletions accelerated net cell proliferation in vitro, consistent with our in vivo results (Fig. 4e). Overall, these findings suggest that the acceleration in tumor recurrence observed for cells harboring deletions *GPX4*, *GOT1*, or *PNPLA2* is unlikely to be due to an increased rate of tumor growth, resulting in a decreased latency to clinical detection.

### Deletion of *PNPLA2*, *GOT1*, or *GPX4* may Increase the Probability of Dormant Tumor Cell Reentry into the Cell Cycle

We next considered a third possibility in which deletion of *PNPLA2*, *GOT1*, or *GPX4* might increase the probability of cell cycle reentry from the dormant state. Prior data from our model suggests that recurrent tumors arise stochastically from rare dormant tumor cells that reenter the cell cycle (11,14,15). As such, for a given number of dormant residual tumor cells, an increased probability per cell per unit time that a cell exits dormancy and reenters the cell cycle would be predicted to accelerate tumor recurrence.

To test this hypothesis, we studied the rate of EdU incorporation in dormant residual lesions as a function of time and of genotype for *GPX4*, *GOT1*, and *PNPLA2*. Primary orthotopic tumors formed from sg*GPX4*, sg*GOT1*, and sg*PNPLA2* tumor cells regressed following Her2 downregulation induced by doxycycline withdrawal. Residual lesions were harvested at days 10 and 28 following doxycycline withdrawal (Fig. 5a). Mice received 3 intraperitoneal injections of EdU: 24, 18, and 2 hr prior to sacrifice. Rates of tumor cell EdU incorporation were measured in tissue sections that were co-stained for GFP to identify tumor cells.

**Figure 5:**
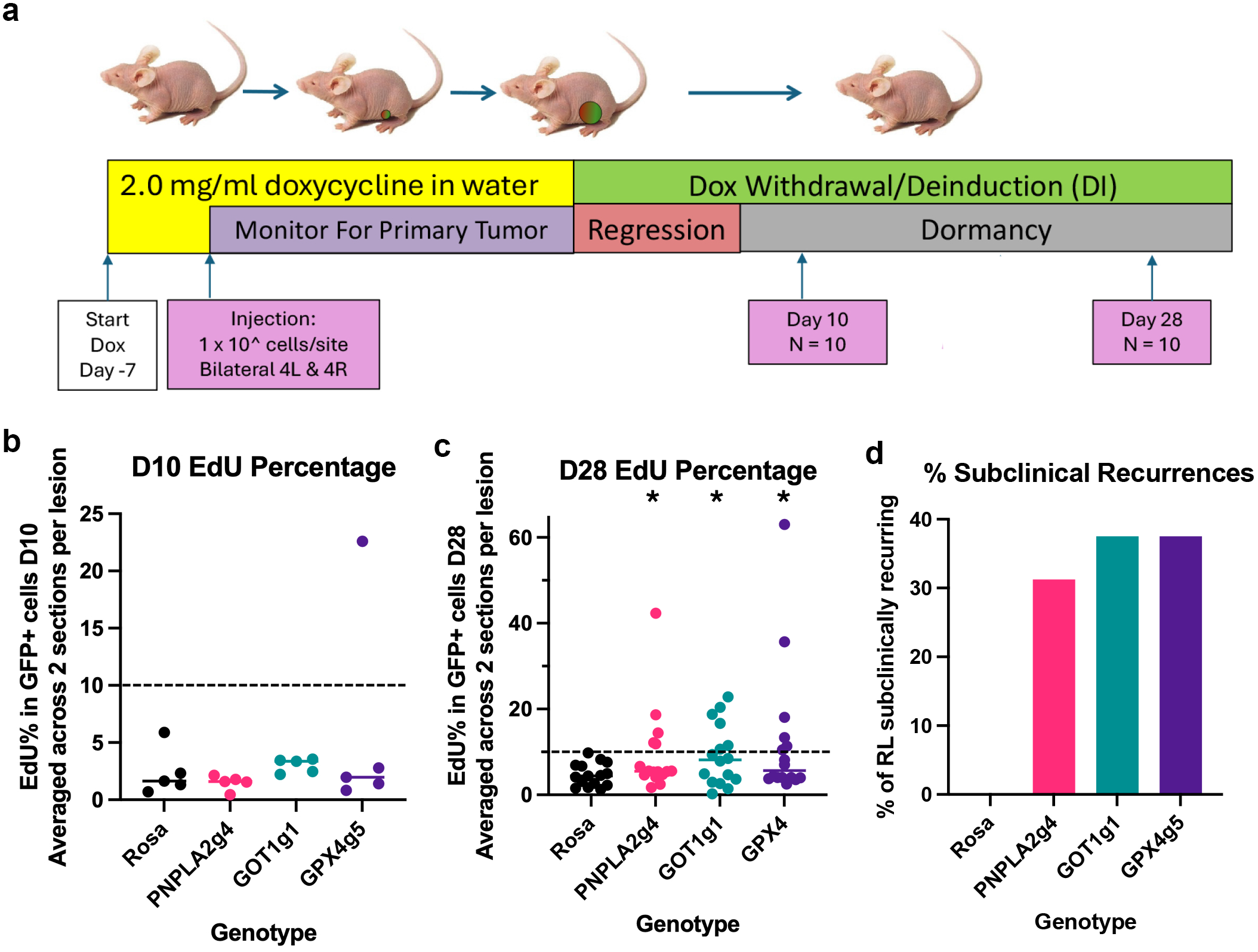
Deletion of ferroptotic regulators increases probability of cells reentering the cell cycle a. Schematic of experiment harvesting residual lesions to assess EdU percentages. b. Day 10 residual lesion EdU percentage. Each point is an average of EdU percentage from two non-adjacent tissue sections. n = 5 for each genotype. c. Day 28 residual lesion EdU percentage. Each point is an average of EdU percentage from two non-adjacent tissue sections. n = 15 for Rosa, n = 16 for *PNPLA2g4*, *GOT1g1*, *GPX4g5*. d. Barplot demonstrating % of total number of lesions examined that were sub-clinically recurring in each genotype.

As anticipated, 24 hr EdU incorporation rates at D10 were extremely low in 18 of 20 samples across all 4 genotypes (range = 0.44%-4.36%, median = 2.08) (Fig. 5b). One of five sg*Rosa* lesions evaluated had a relatively low rate of EdU incorporation that was nonetheless an outlier (7.64%). Surprisingly, one of five sg*GPX4* lesions tested was found to be already subclinically recurring at D10 (EdU = 22.6%). Thus, while there was some variability across groups, the low EdU percentages observed is consistent with the inference that the majority of residual lesions were dormant across all genotypes tested.

At day 28 following doxycycline withdrawal, all 15 sg*Rosa* lesions examined exhibited 24 hr EdU incorporation rates <10%, suggesting that these lesions remained in a dormant state at this timepoint (Fig. 5c). In contrast, all 3 gene deletion groups exhibited bimodal distributions wherein one set of residual lesions exhibited low 24 hr EdU incorporation rates similar to those observed for sg*Rosa* cells, and another set of residual lesions exhibited substantially higher 24 hr EdU incorporation rates than those observed for sg*Rosa* cells. These elevated EdU incorporation rates suggest that one or more cells in these lesions had exited the dormant state, reentered the cell cycle, and given rise to a detectable population of proliferating cells.

We used a threshold of 10% EdU incorporation to classify residual lesions as dormant vs. subclinical recurrences, since a threshold of 10% approximately bisected the bimodal EdU-low and EdU-high residual lesions. This resulted in the finding that none of 15 sg*Rosa* residual lesions had reactivated, whereas 5 of 16 sg*PNPLA2* lesions, 6 of 16 sg*GOT1* lesions, and 6 of 16 sg*GPX4* lesions had reactivated (Fisher’s exact test: p = 0.043, sg*PNPLA2* vs. sg*Rosa*; p = 0.018, sg*GOT1* vs. sg*Rosa*; and p = 0.018, sg*GPX4* vs. sg*Rosa,* Fig. 5d). This finding is consistent with a model in which dormant tumor cells with deletion of *PNPLA2*, *GOT1*, or *GPX4* exhibit an increased probability of reentering the cell cycle as a function of time.

### Dormant Cells are Resistant to Ferroptosis Inducers Erastin and RSL3

To examine whether the effects of PNPLA2, GOT1, and GPX4 on dormancy and tumor recurrence were mediated by ferroptosis, we studied the effects of ferroptosis inducing agents in dormant and proliferating cells in vitro. First, proliferating Cas9-expressing iHer2 cells maintained under full growth conditions in the presence of doxycycline were treated with the ferroptosis-inducing agent erastin, which selectively and irreversibly inhibits the cystine-glutamate antiporter, System Xc^-^ (17,40,41).

Proliferating Cas9-expressing iHer2 cells were treated with increasing doses of erastin from 10nM to 100µM for 24, 48, or 72 hr and viable cell number was measured using an MTT assay (42,43). At 24 hr, a decreased number of viable tumor cells was evident following 1µM erastin treatment, with 10µM treatment resulting in a ∼50% reduction in viable tumor cell number compared with vehicle-treated cells (Fig. 6a). Increasing durations of erastin treatment in the range of 1-100µM resulted in incrementally greater decreases in viable cell number, although the majority of cell death occurred within 24 hr (Supp. Fig. 4a, b).

**Figure 6:**
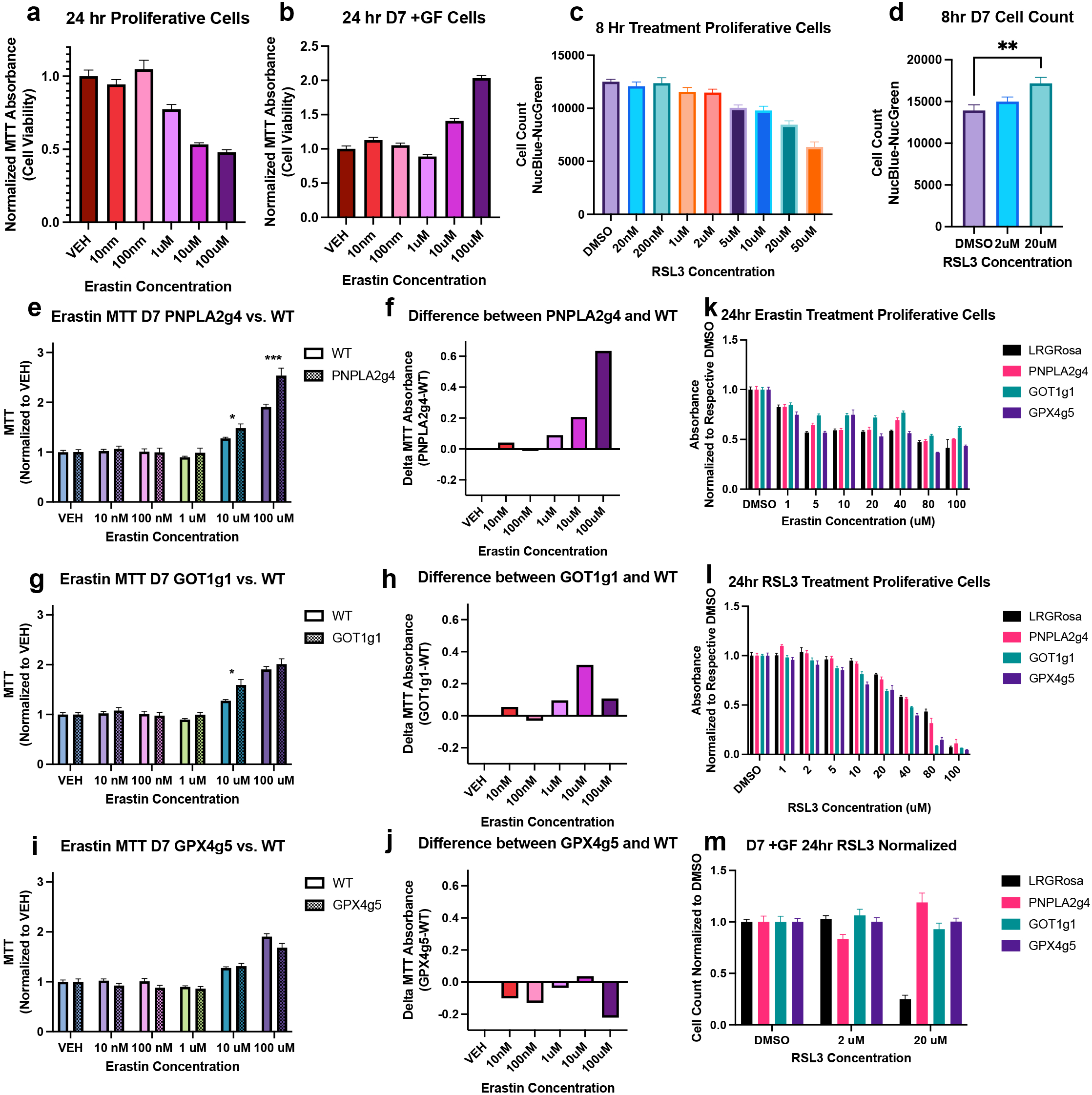
***Dormant residual tumor cells exhibit resistance to ferroptosis inducers in vitro*** Results of MTT assay after treating: a. proliferative iHer2 cells, b. iHer2 cells at D7 of dormancy with increasing doses of erastin for 24hr. MTT absorbance in each experiment, as a proxy for cell viability, is normalized to its own vehicle control (equal volume DMSO). Cell counts after treating: c. proliferative iHer2 cells and d. iHer2 cells at D7 of dormancy with increasing doses of RSL3 for 8hr. Cells were imaged by EVOS m5000 and live cells were quantified by NucBlue staining for all nuclei and NucGreen staining for dead nuclei. e-j. iHer2 cells with KO of gene of interest (*PNPLA2*, e-f, *GOT1*, g-h, *GPX4*, i-j) at D7 of dormancy were treated with erastin for 24 hrs and the results of MTT assay at 24hr are shown. e, g, and i display normalized MTT absorbance as compared to wildtype iHer2 cells, with f, h, and j displaying the difference between the KO and WT at each dose. k, l. Results of MTT assay after treating proliferative iHer2 cells with knockouts of *PNPLA2*, *GPX4*, *GOT1*, or *LRGRosa* with increasing doses of erastin (k) or RSL3 (l) for 24hr. m. Cell counts after treating iHer2 cells with knockouts of *PNPLA2*, *GPX4*, *GOT1*, or *LRGRosa* at D7 of dormancy with RSL3 for 24 hours. Cells were imaged on EVOS as above.

Having demonstrated the anticipated effects of erastin on proliferating iHer2 cells, we next tested the effects of erastin on iHer2 cells at day 7 of dormancy. Strikingly, while proliferating iHer2 cells were erastin-sensitive, dormant iHer2 cells were resistant to erastin treatment (Fig. 6b). Indeed, rather than the anticipated decrease in viable cell number in response to erastin treatment, the number of viable dormant tumor cells was unchanged by treatment with 10nM-1µM erastin for 24-72 hrs, and actually increased after treatment with 10µM or 100µM erastin for 24-72 hr during the dormancy window (Supplementary Fig. 4c-f).

To confirm that these findings accurately reflected changes in viable cell number, we repeated the above in vitro dormancy experiments using microscopic imaging to directly measure cell number. Following drug treatment, 5% NucBlue dye – which is similar to Hoechst 33342 (44) – was added to the media in each well, incubated for one hr and imaged. QuPath analysis software was then used to count the total number of nuclei in each well based on captured images.

This approach confirmed the erastin-resistant phenotype observed in dormant tumor cells using the MTT assay. Specifically, treatment of proliferating iHer2 cells with increasing doses of erastin resulted in the anticipated decreases in cell number, whereas under dormancy conditions tumor cell number was unchanged by treatment with 10nM-1µM erastin and actually increased following treatment with 10-100µM erastin (Supp. Fig 5a, b).

After demonstrating the counterintuitive effects of erastin on dormant tumor cells, we wished to address whether this phenotype was erastin-specific. Accordingly, we examined the response of proliferative vs. dormant cells in response to a second ferroptosis inducer, RSL3, which inhibits GPX4 (35). Recent studies have called into question RSL3’s mechanism of action and suggest that RSL3 may bind not GPX4, but a different selenoprotein, TXNRD1, that also has reducing capabilities in the cell (45–47).

Following 8 hr of RSL3 treatment, reduced viable cell numbers were observed for proliferating cells treated with doses of RSL3 ≥ 2 µM compared with DMSO treated controls (p = 0.027) (Fig. 6c). In contrast, treating D7 dormant tumor cells with RSL3 for 8 hr failed to decrease cell number. In fact, treatment with 20µM RSL3 for 8 hr resulted in a statistically significant increase in cell number in dormant tumor cells compared to DMSO control (Fig. 6d). This finding parallels our findings with erastin, suggesting that the resistance to ferroptosis that we observe in dormant tumor cells may constitute a generalized response, rather than an erastin-specific effect.

### *PNPLA2*, *GOT1*, and *GPX4* Deletion Modulate Sensitivity to Erastin and RSL3

To determine whether GPX4, GOT1, or PNPLA2 influence the sensitivity of tumor cells to agents that induce ferroptosis, we performed MTT assays following erastin treatment of cells in which *GPX4*, *GOT1*, or *PNPLA2* had been deleted. Cells with *PNPLA2* knockout (KO) exhibited an exaggerated response to erastin treatment during dormancy. Specifically, at higher erastin doses at which dormant iHer2 cells exhibited increased viable tumor cell number, this counterintuitive effect was further enhanced in cells with loss of *PNPLA2* (Fig. 6e, f). If the increase in viable tumor cell number observed following high levels of erastin treatment reflects the reentry of dormant tumor cells into the cell cycle, this finding suggests that dormant *PNPLA2* KO cells may be more likely to reenter the cell cycle, which would be consistent with the increased rate at which residual lesions composed of *PNPLA2* KO cells spontaneously reactivate and recur in vivo.

Dormant *GOT1* KO cells exhibited a more modest enhancement of the counterintuitive effect of erastin treatment compared to *PNPLA2* KO cells, but nevertheless displayed a trending increase in viable tumor cell number at 10µM erastin as assessed by MTT (Fig. 6g, h). Dormant *GPX4* KO cells did not exhibit an enhanced counterintuitive response to high dose erastin treatment and instead trended towards decreased resistance to erastin at 100µM (Fig. 6i, j).

Our findings confirmed that dormant tumor cells exhibit resistance to both erastin-induced and RSL3-induced cell death, as well as a counterintuitive increase in viable tumor cell number following treatment with high doses of either of these agents. Moreover, loss of *PNPLA2* and, to a lesser extent, *GOT1* exaggerated the counterintuitive effects observed at high doses of erastin. Thus, resistance to erastin and, by inference, ferroptosis appears to be a consistent feature associated with the dormant state.

In light of evidence supporting their ferroptosis-suppressing roles, we anticipated that deletion of *PNPLA2*, *GOT1* or *GPX4* would promote ferroptotic cell death under proliferative conditions, and thus lead to further decreases in cell survival following treatment with erastin. In contrast, *PNPLA2* KO did not significantly alter erastin response at 24 hr of treatment under proliferative conditions compared to *Rosa* control (Fig. 6k). Similarly, *GPX4* deletion had little, if any, effect on erastin sensitivity under proliferative conditions. Surprisingly, *GOT1* KO provided a modest but consistent protective benefit at doses of erastin between 5µM and 40µM (5µM, p=6E-6; 10µM, p=8.2E-5; 20µM, p=8.2E-5; 40µM, p=6E-6).

We next extended our studies of RSL3 findings to cells deleted for *PNPLA2, GOT1* or *GPX4*. As with erastin, we predicted that deletion of these genes would result in increased susceptibility to ferroptotic cell death under proliferative conditions, based on their reported ferroptosis-suppressing functions. Whereas *PNPLA2* KO cells were not significantly different than sg*Rosa* control, both *GOT1* KO and *GPX4* KO cells exhibited significantly increased sensitivity to RSL3 treatment compared to sg*Rosa* controls at RSL3 doses from 10-80µM (Fig. 6l). This suggests that proliferating cells deleted for *PNPLA2, GOT1* or *GPX4* are more susceptible to ferroptosis.

With a longer 24 hr RSL3 treatment as opposed to 8 hr treatment, cell number decreased with 20µM RSL3 treatment in D7 dormant sg*Rosa* cells (Fig. 6m). In contrast, following deletion of *PNPLA2*, *GOT1*, or *GPX4*, D7 dormant cells were each highly resistant to 24 hr of treatment with 20µM RSL3 (Fig. 6m).

In summary, *PNPLA2* KO enhanced the counterintuitive increase in viable cell number observed in response to high-dose erastin treatment, but only in the dormant state. *GOT1* KO exhibited a more modest enhancement of this counterintuitive effect in the dormant state, as well as a decrease in erastin sensitivity under proliferative conditions, whereas *GPX4* KO did not influence erastin response. Deletion of *GOT1* or *GPX4* resulted in increased RSL3 sensitivity under proliferative conditions, whereas deletion of *PNPLA2*, *GOT1*, or *GPX4* promoted marked resistance to RSL3-mediated ferroptosis during dormancy. Taken together, these findings indicate that dormancy is a ferroptosis-resistant state, and that resistance is at least partially augmented by deletion of *PNPLA2*, *GOT1*, or *GPX4*.

## Erastin and RSL3 Sensitivity of iHer2 Cells is Partially Reversed by Ferrostatin-1

Though erastin is considered to function as a canonical inducer of ferroptosis, in certain contexts it has also been reported to induce apoptosis (48). Thus, we wished to confirm that the observed effects of erastin in our experiments were mediated by ferroptosis. One approach to evaluate this would be to demonstrate that erastin’s effects can be reversed by treatment with a ferroptosis inhibitor, such as ferrostatin-1. In this regard, treatment of cells with erastin or RSL3 in combination with 100uM ferrostatin-1 rescued the effects of these ferroptosis-inducing agents on viable cell number to levels similar to DMSO control, whereas treatment with 10µM ferrostatin-1 yielded lower levels of rescue (Supp. Fig. 6a-c). An increase in viable cell number was observed at lower doses of ferrostatin-1 by pretreating proliferating cells with ferrostatin for 24 hr prior to treatment with the ferroptosis inducing agent (Supp. Fig. 6d). In this treatment paradigm, treatment with either 1 µM or 10µM of ferrostatin-1 for 24 hr prior to treatment with 20µM RSL3 significantly increased estimated viable cell number (RSL3 vs. RSL3 + 1uM FS, p = 0.0019; RSL3 vs. RSL3 + 10uM FS, p = 0.0061).

Under dormancy conditions, ferrostatin-1 treatment in combination with erastin further increased estimated viable cell number, even under conditions where erastin itself increased viable cell number (Supp. Fig 6e). This finding provides additional evidence supporting the resistance to erastin exhibited by dormant tumor cells, but indicates that the mechanism by which erastin increases viable cell number is not reversible by ferrostatin-1 treatment, thereby suggesting that it is likely not directly attributable to effects on lipid peroxidation. In the case of cells treated with either 10µM or 100µM RSL3, addition of 100µM ferrostatin was able to partially rescue cell viability (Supp. Fig. 6f). In summary, we found that erastin and RSL3-induced cell death was at least partially reversible with ferrostatin-1, leaving open the possibility that processes other than ferroptosis may contribute to stark differences in sensitivity to ferroptosis induction in dormant vs. proliferating cells.

## Glutathione and Adaptation to Lipid Peroxides May Contribute to Ferroptosis Resistance in Dormant Cells

In attempting to understand the observed resistance to ferroptosis exhibited by dormant tumor cells, we hypothesized that the cellular environment of dormant cells may lack a critical component required for ferroptotic cell death. In particular, the reduced form of glutathione (GSH) constitutes a key protection against ferroptotic cell death. Accordingly, a relative lack of GSH within the cell promotes ferroptotic death. Therefore, we measured the ratio of reduced to oxidized glutathione (GSH/GSSG) in proliferating and dormant cells. In sg*Rosa* control cells, D14 dormant tumor cells exhibited a 2.3-fold increase in GSH/GSSG ratio compared to proliferative cells, indicative of a more highly reduced cellular environment (Fig. 7a).

**Figure 7:**
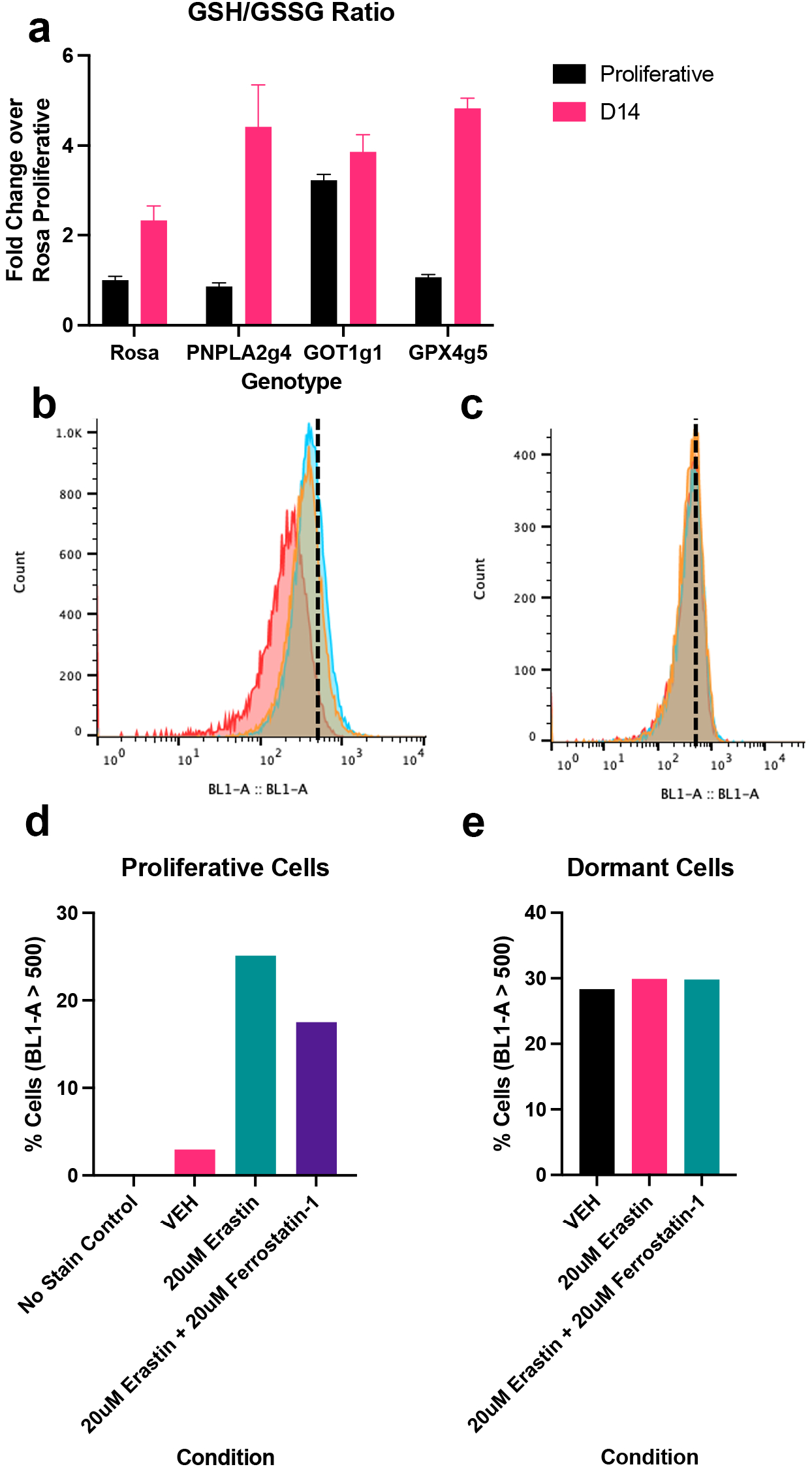
Glutathione and lipid oxidation measurements of dormant tumor cells consistent with ferroptosis resistance a. Reduced to oxidized glutathione ratio in proliferating and dormant (D14) sg*Rosa*, sg*PNPLA2*, sg*GOT1*, and sg*GPX4* cells. Data presented as fold-change over proliferative sg*Rosa* GSH/GSSG ratio b. Distribution of fluorescence of proliferative untransduced iHer2 cells stained with Liperfluo following treatment with VEH (red), 20 µM erastin (blue), or 20 µM erastin and 20 µM ferrostatin (orange) for 24 hr. c. Distribution of fluorescence of dormant D10 untransduced iHer2 cells stained with Liperfluo following treatment with VEH (red), 20 µM erastin (blue), or 20 µM erastin and 20 µM ferrostatin (orange) for 24hr. Dashed vertical line is placed at BL1-A =500. d, e. Barplot displaying percentage of proliferative (d) or dormant (e) cells stained with Liperfluo with BL1-A measurements > 500.

Under proliferative conditions, *PNPLA2* and *GPX4* KO cells exhibited a GSH/GSSG ratio similar to that in sg*Rosa* control cells. Intriguingly, proliferating *GOT1* KO cells displayed a ∼3- fold increase in GSH/GSSG compared to sg*Rosa* cells, which is consistent with our prior observation that proliferating *GOT1* KO cells exhibit decreased sensitivity to erastin.

At D14 of dormancy, *PNPLA2* KO, *GOT1* KO, and *GPX4* KO cells each exhibited an increased GSH/GSSG ratio compared to their proliferative state. GSH/GSSG ratios for dormant *PNPLA2*, *GOT1* and *GPX4* KO cells were each approximately 4-fold higher than the proliferative sg*Rosa* controls and ∼2-fold higher than the D14 dormant sg*Rosa* control (Fig. 7a).

Together, these results indicate that dormant iHer2 cells have a higher GSH/GSSG ratio, which would be anticipated to result in increased resistance to ferroptotic cell death due to oxidative stress. Furthermore, our observation that deletion of *PNPLA2*, *GOT1* or *GPX4* each result in greater increases in the GSH/GSSG ratio compared to Rosa controls in the dormant state suggests a potential mechanism by which deletion of *PNPLA2*, *GOT1* or *GPX4* could increase resistance to erastin.

Oxidized lipids are a direct effector of ferroptotic cell death (49). Accordingly, differences in lipid oxidation levels between dormant and proliferative cells could potentially explain their differential sensitivity to ferroptotic death. In order to measure oxidized lipids, we used Liperfluo, a fluorescent probe for lipid peroxides that emits at 535nm when oxidized (50). We examined both proliferative and D10 dormant cells treated for 24 hr with erastin, erastin plus ferrostatin, or vehicle control.

In proliferating cells, ∼3% of vehicle-treated cells exhibited lipid peroxidation by comparing fluorescence in stained vs. unstained cells (Fig. 7b, d). Following treatment with 20µM erastin, 25% of cells exhibited lipid peroxidation. Treatment of proliferating cells with ferrostatin, a lipophilic antioxidant, concordantly with erastin resulted in the partial rescue of this erastin-induced increase, as 17.5% of cells were lipid peroxide-positive (Fig. 7d).

The lipid peroxide profile of dormant tumor cells was markedly different from that of proliferating cells. Using the same fluorescence threshold as used for proliferative cells, 28% of vehicle-treated dormant cells were considered lipid peroxide-positive (Fig. 7c, e). Notably, treatment of dormant cells with erastin or erastin plus ferrostatin did not materially change this distribution, with approximately 30% of cells being identified as positive under both treatment conditions. This suggests the possibility that the erastin-resistant phenotype that we observe in dormant tumor cells may be partially explained by an inability of erastin to further increase lipid peroxidation. However, our observation that lipid peroxidation in vehicle-treated cells is substantially higher in dormant compared to proliferating cells suggests the possibility that the dormant state may have additional adaptations to survive lipid peroxidation and/or mitigate the effects of additional lipid peroxidation that could otherwise result in ferroptotic cell death.

## Loss of *PNPLA2*, *GOT1*, or *GPX4* is Associated with Worse Outcomes in Human Breast Cancer Patients

To begin to evaluate the potential clinical relevance of our findings, we examined expression and copy number alterations in *PNPLA2*, *GOT1*, and *GPX4* in relation to breast cancer outcomes. We examined data for > 2,000 breast cancer patients from METABRIC (Molecular Taxonomy of Breast Cancer International Consortium), including long-term patient outcomes. Consistent with our results in mice, loss of either *PNPLA2* or *GPX4* in primary breast cancers was significantly associated with poorer recurrence-free survival outcomes in breast cancer patients across all tumor subtypes (*PNPLA2*: HR=1.3, p=0.0077; *GPX4*: HR=1.2, p=0.026) (Fig 8a, c). Although *GOT1* loss was not significantly associated with recurrence outcomes across all tumor subtypes (Fig 8b), when limited to hormone receptor (HR)-positive tumors, loss of each gene was significantly associated with poorer recurrence-free survival (*PNPLA2*: HR=1.5, p=2.3e-4; *GOT1* HR=1.4, p=0.013; *GPX4* HR=1.3, p=0.014) (Fig. 8d-f).

**Figure 8:**
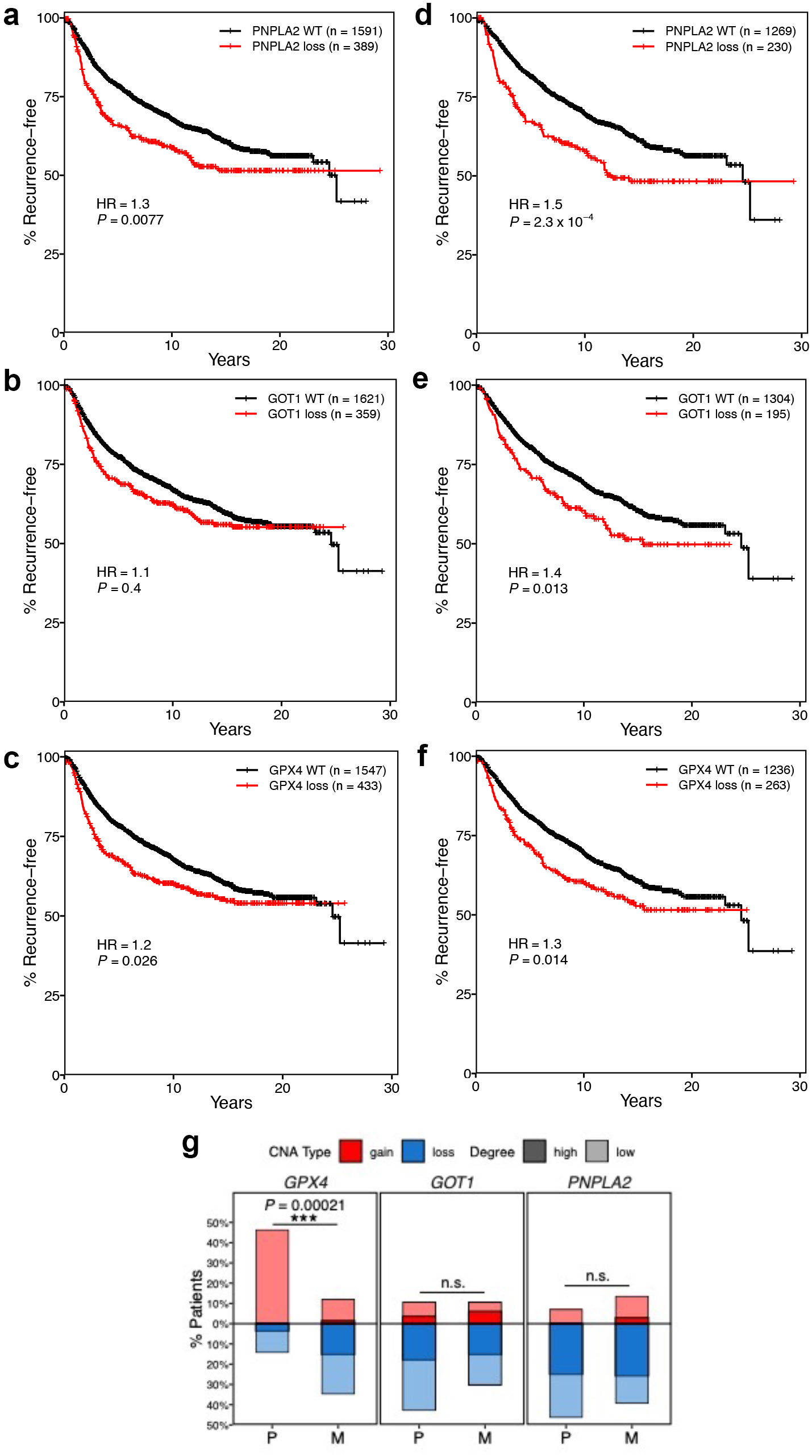
Human breast cancer genomic and transcriptomic data supports role of PNPLA2, GOT1, and GPX4 loss in accelerating recurrence a-c. Recurrence-free survival outcomes for METABRIC patients harboring loss of *PNPLA2* (a), *GOT1* (b), and *GPX4* (c) across all tumors. d-f. Recurrence-free survival outcomes for METABRIC patients harboring loss of *PNPLA2* (d), *GOT1* (e), and *GPX4* (f) restricted to HR+ tumors. g. Percentage of patients in METAMORPH (56) study harboring copy number changes in *PNPLA2*, *GOT1*, or *GPX4* in primary or metastatic tumors.

A similarly significant pattern of associations across all tumors was evident when examining overall survival as opposed to recurrence-free survival, and this association was more pronounced among HR-positive tumors, (Supp. Fig. 7a-f). Further analysis of genomic data revealed that copy loss was highly prevalent in metastatic tumors for each of these three genes (Fig. 8g). Copy number loss for *PNPLA2* and *GOT1* was also prevalent in primary tumors, but a significant increase in copy number loss in metastases compared to primary tumors was evident for *GPX4*. Altogether, the associations of CN loss with poorer survival outcomes in breast cancer patients are generally concordant with the results of our in vivo studies in mice.

Similar associations emerged when examining gene expression as opposed to copy number alterations. *PNPLA2* and *GPX4* expression were highest amongst ER+ and low-grade primary breast cancers, which typically have the best outcomes (Supp. Fig. 8a, b (*PNPLA2*), 8e, f (*GPX4*)). *GOT1* expression did not follow this pattern and was instead expressed at highest levels in ER-negative and high-grade tumors, which may partially explain why GOT1 loss was only associated with better outcomes when the analysis was restricted to HR+ tumors (Supp. Fig. 8c, d). Furthermore, when examining recurrence outcomes, expression of *PNPLA2* and *GPX4* were each associated with better 5-year relapse-free survival, as anticipated from our recurrence-free survival analysis in mice (Supp. Fig. 9a, c). *GOT1* exhibited the opposite association, perhaps due to its association with different grade and subtype of tumors (Supp. Fig. 9b).

## Discussion

Our findings presented here identify two counterintuitive effects of *PNPLA2, GOT1,* and *GPX4* deletion on ferroptosis sensitivity and tumor recurrence. First, our data reveal that deleting *PNPLA2, GOT1,* or *GPX4* each accelerate tumor recurrence in genetically engineered mouse models, in contrast to expectations based on their ferroptosis-suppressing functions.

Second, our data reveal that dormant tumor cells exhibit intrinsic resistance to ferroptosis, and that deletion of *PNPLA2, GOT1,* or *GPX4* accentuates the resistance of dormant tumor cells to ferroptotic stimuli. Consistent with the above observations, we found evidence that primary breast cancers in patients that express decreased levels of these genes, or have undergone copy number loss, exhibit accelerated recurrence.

Notably, our findings indicated that deletion of *PNPLA2, GOT1,* or *GPX4* did not alter tumor cell fitness, either during acute tumor regression in response to Her2 downregulation or during the dormant period, and did not accelerate recurrence by virtue of increasing the rate of tumor cell proliferation in recurrent tumors. Instead, we found evidence suggesting that deletion of each of these genes accelerates tumor recurrence at the cellular level by promoting dormant tumor cells to re-enter the cell cycle, thereby initiating tumor recurrence sooner than would otherwise have occurred. Together, these data provide a cohesive model explaining our counterintuitive findings that deletion of *PNPLA2, GOT1,* or *GPX4* each accelerates tumor recurrence.

Our findings provide evidence that dormancy is a ferroptosis-resistant cell state. As anticipated, treatment of proliferating iHer2 cells with increasing doses of the ferroptosis inducers erastin or RSL3 resulted in a decrease in the number of viable tumor cells, as has been repeatedly described in the literature (51). In contrast, in response to increasing doses of erastin or RSL3, the number of viable dormant tumor cells either remained unchanged or increased. Moreover, we found that CRISPR-Cas9 deletion of *PNPLA2* or *GOT1* enhanced dormant tumor cells resistance to both erastin and RSL3, whereas *GPX4* deletion enhanced resistance to RSL3, but not erastin, in the dormant state. Thus, while proliferating iHer2 cells reproducibly exhibit ferroptotic cell death in response to erastin and RSL3, these same cells become highly resistant to these agents when dormant.

While the acceleration in tumor recurrence that we observed in mice following deletion of *PNPLA2*, *GOT1*, or *GPX4* could be mediated by ferroptosis resistance in dormant tumor cells, these could also be mechanistically unrelated phenotypes. By promoting residual tumor cell survival under conditions of oxidative stress, resistance to a mechanism of cell death would be anticipated to result in a greater number of surviving dormant residual tumor cells. This would, in turn, give rise to a larger pool of residual tumor cells from which a stochastic recurrence could arise. Importantly, such a mechanism would also be anticipated to preferentially select for cells deleted for *PNPLA2*, *GOT1* or *GPX4* in either in vitro or in vivo competition assays. In contrast to this prediction, we did not observe selection for or against tumor cells harboring guides targeting these genes. This suggests that the acceleration in tumor recurrence that we observed in our models was unlikely to be due to a competitive survival advantage manifested by cells deleted for these genes, either immediately following therapy or when dormant. Rather, our data point to an effect of *PNPLA2*, *GOT1* or *GPX4* deletion that occurs later in process of recurrence, whereby dormant tumor cells deleted for *PNPLA2*, *GOT1* or *GPX4* exhibit an increased probability of re-entering the cell cycle to give rise to a recurrent tumor.

Nevertheless, we note that if the enhanced survival of dormant tumor cells deleted for *PNPLA2*, *GOT1* or *GPX4* is due to a non-autonomous or paracrine effect, this would theoretically result in a greater number of surviving dormant tumor cells without changing the ratio of deleted:wildtype cell populations. In this regard, such a model would be consistent with the recurrence-free survival curves that we observed in our in vivo cell competition assays. In particular, given the fact that only ∼20% of injected cells harbored guides targeting *PNPLA2, GOT1*, or *GPX4* in these experiments, the acceleration in recurrence that we observed compared to the sg*Rosa*:sg*Rosa* control was unexpected. However, if there was an overall increase in the population of surviving residual tumor cells as a whole, this larger pool of cells from which a stochastic recurrence could emerge would be compatible with the observed acceleration in tumor recurrence. Consequently, further studies would be of value to evaluate changes in the number of surviving dormant tumor cells, rather than simply the ratio of mutant:wildtype cells. Nevertheless, differences in cell survival for cells harboring deletions in *PNPLA2*, *GOT1* or *GPX4* were not observed in vitro, arguing against the paracrine model.

An alternative possibility is that the accelerated tumor recurrence that we observed in cells deleted for *PNPLA2*, *GOT1* or *GPX4* is mediated by ferroptosis resistance via a mechanism related to the phenotype that we describe in which the number of viable tumor cells increased following treatment of dormant tumor cells with higher concentrations of ferroptosis inducing agents. If these gene deletions, which based on the literature would be anticipated to be ferroptosis-promoting, act on dormant cells in an analogous manner to pharmacological ferroptosis inducing agents, such as erastin and RSL3, an intriguing possibility is that each may act to increase the likelihood of dormant cell reentry into the cell cycle. This is consistent with our EdU incorporation results in D28 residual lesions suggesting that cells deleted for *PNPLA2*, *GOT1* or *GPX4* are more likely to reenter the cell cycle, though the mechanism for such an effect would require further study.

We hypothesized that the resistance to ferroptosis observed in dormant tumor cells might be due to an altered cellular state lacking one or more components or conditions required for ferroptotic death, including reduced antioxidant capacity, lipid oxidation, or iron. In this regard, the markedly increased GSH/GSSG ratio that we detected in dormant tumor cells compared to proliferative tumor cells, which was accentuated in cells harboring sg*PNPLA2,* sg*GOT1,* or sg*GPX4*, could potentially constitute one such lacking component, since these increased levels of glutathione may enable the suppression of lipid peroxidation below the threshold required for death to occur.

In contrast to our findings of increased GSH/GSSG ratio and resistance to erastin and RSL3-induced cell death in iHer2 cells harboring sg*GOT1*, *GOT1* knockdown has previously been reported to decrease the GSH/GSSG ratio and also to sensitize cells to ferroptotic cell death (34,52). However, *GOT1* dependence has been reported to be tissue-specific, which could explain the present result (53). In particular, *GOT1* knockout dramatically altered cellular metabolism and slowed growth in pancreatic cancer cells, but barely impacted colorectal cancer cell metabolism or growth. *GOT1* deletion may promote glutamate transamination to form phosphoserine, which can then contribute to glycine production (54). Glycine is one of the three amino acids needed for reduced glutathione production (55). Thus, *GOT1* deletion may alter downstream metabolism to increase the reduced form of GSH, which would ostensibly prevent ferroptosis by increasing the GSH/GSSG ratio.

In summary, we observed counterintuitive effects of deleting *PNPLA2*, *GOT1* or *GPX4* on accelerating tumor recurrence. As these three genes are each predicted to have anti-ferroptotic functions, inhibiting ferroptosis would reasonably be considered to constitute a viable therapeutic target to induce cancer cell death. In contrast to this expectation, our results suggest that targeting these genes might have the potential to decrease clinical latency and accelerate tumor recurrence by promoting the re-entry of dormant residual tumor cells into the cell cycle. Thus, caution may be warranted in pursuing such a therapeutic strategy, and further studies are needed to better understand the underlying mechanisms.

Further, our results demonstrate that dormant cells are intrinsically ferroptosis resistant. Lee et. al. recently demonstrated that cell cycle arrest can promote ferroptosis resistance (23), whereas Puente-Cobacho et. al. demonstrated that dormant D2.OR cells are intrinsically ferroptosis resistant (24). Our studies are consistent with these findings, but extend their implications by showing that therapy-associated cancer cell dormancy is associated with profound ferroptosis resistance, rather than chemically-induced cell cycle arrest or microenvironment-induced dormancy. Additionally, we implicate three ferroptosis-modulating genes in this resistance phenotype by demonstrating counterintuitive effects of *PNPLA2*, *GOT1* or *GPX4* deletion on tumor recurrence in a model in which cells pass through a dormant state, and provide a potential mechanism of accelerated rates of dormancy exit as a contributor to recurrence. Together with prior work in this area, our findings contribute to the broader understanding of a mechanism of cancer cell adaptation that may materially contribute to mortality from breast cancer, and thus constitute a potential therapeutic target.

## Methods

### In vitro assays

MTB/TAN-derived primary tumor cells were cultured in DMEM (Corning, Cat. # 10-017- CV) containing 10% bovine calf serum supplemented (Cytiva Hyclone, Cat. # SH30072), 1% Penicillin/Streptomycin (Thermo Fisher Scientific, Cat. # 15-140-122), 1% Glutamine (Thermo Fisher Scientific, Cat. # 25030081), 2mg/ml Doxycycline (RPI, Cat. #D43020- 250.0), 5mg/ml Insulin (GeminiBio, Cat. # 700–112P), 10ug/ml EGF (Millipore, Cat. # E4127), 1mg/ml Hydrocortisone (Sigma, Cat. # H0396), 1mM Progesterone (Sigma, Cat. # P7556).

For in vitro dormancy experiments, cells were plated on D-3 in medium as above, transitioned to medium as above containing 1% bovine calf serum on D-2, and transitioned to medium containing 1% bovine calf serum and no Doxycycline on D0. Plates were harvested at dormancy time points as desired, while medium containing 1% bovine calf serum and no doxycycline was replaced on remaining plates weekly. Reinduction plates were harvested at time points as desired following 48h treatment with medium containing 1% bovine calf serum and doxycycline as indicated in the figures.

### Animal experiments

All animal studies were approved by the University of Pennsylvania Institutional Animal Care and Use Committee (IACUC). In vivo competition assays and recurrence-free survival assays were performed as previously described. Briefly, 1e06 cells (1:1 mixture of GFP^+^ and mCherry^+^ cells for some competition assays or unmixed cells for recurrence-free survival assays and the CRISPR-Cas9 screen) were transplanted into the inguinal #4 mammary fat pads of nu/nu female mice (NCRNU-F, Taconic) and administered 2mg/ml doxycycline/5% sucrose via drinking water. One competition assay injected 8e05 cells and an 20:80 mixture of GFP^+^ to mCherry^+^ cells, with otherwise identical methodology. Mice were de-induced by switching to regular drinking water once mammary tumors reached a target size (5×5mm for competition assays and recurrence-free survival assays, and 3x3mm for the CRISPR-Cas9 screen). Mice were palpated two or three times per week and time-to-recurrence was assessed by Kaplan-Meier analysis as measured by when non-palpable, dormant lesions first became palpable, or when stably regressed lesions began to increase in size. Hazard ratios and p-values were calculated using the logrank test. At the time of harvest, mice were administered 50mg/kg EdU (i.p.) 2h prior to euthanasia, or in the case of the residual lesion experiment, 50 mg/kg EdU was administered 24h, 18h, and 2h prior to euthanasia.

Primary and recurrent tumor volume data from animal models were converted to tumor size estimates using tumor size = (volume × 6 ÷ π)^1/3^ (i.e. the diameter of a spherical object of equivalent volume). Tumor growth rate was defined as change in tumor size as a linear function of time and was estimated using linear regression between tumor size and time. Tumor growth rates between different experimental groups were compared using Wilcoxon rank-sum test.

### Plasmids and lentiviral production

LentiV_Cas9_puro (Addgene, Cat. # 108100) was used to generate Cas9 expressing *MTB/TAN*-derived primary tumor cells. LRG2.1 (Addgene, Cat. # 108098), LRG (Addgene, Cat. # 65656), or LRmCherry2.1 (Addgene, Cat. # 108099) vector backbones were utilized for cloning sgRNAs for CRISPR-Cas9 studies (25). For each sgRNA, sense and antisense oligos were phosphorylated and annealed and then ligated into BsmB1-digested vector. Ligated vectors were transformed into the chemically competent Stbl3 bacteria (Thermo Fisher Scientific, Cat. # C737303). Successfully transformed bacterial clones were picked from Ampicillin selective plates and isolated DNA was sequenced using a U6 primer to confirm sgRNA incorporation.

Lentiviruses were generated in HEK293T cells using the TransIT-293 transfection reagent (Mirus, Cat. # MIR2700) to introduce packaging plasmids pMD2.G (Addgene, Cat. # 12259; 3μg) and psPAX2 (Addgene, Cat. # 12260; 6μg), and 9μg of the desired backbone containing the sgRNA of interest. sgRNA lentiviruses were titered by serial dilution in *MTB/TAN*- derived primary tumor cells using the fluorophore associated with the vector backbones as a readout by on the Attune NxT flow cytometry (Thermo Fisher).

## PCR and ICE analysis

To validate sgRNA editing efficiency, genomic DNA was isolated from Her2-dependent Cas9 unedited and edited cells one week after transduction with sgRNAs (Qiagen, QIAamp DNA mini kit, Cat. # 51304). PCR amplification was performed (98°C 5min → (98°C 5s, <u>T_m_</u> 5s, 72°C 15s) ×30 → 72°C 3min → 4°C) using the Tm calculated for the designed primers used by the online ThermoFisher Scientific Tm calculator. The resulting amplicons were Sanger sequenced and ICE analysis was performed on the sequences to determine editing efficiency using https://ice.synthego.com/<u>#/</u>.

## CRISPR-Cas9 Screen

54 genes were selected for CRISPR-Cas9 screening in vivo based on the ferroptosis literature at the time of library design. In addition, positive control sgRNAs targeting genes that are known to be pro-proliferative and pro-survival and negative control sgRNAs that are non-targeting were also included in this library. For the 54 genes, 5 sgRNAs were designed for each gene resulting in a total of 331 sgRNAs when including controls. Additionally, a negative control library of non-targeting sgRNAs was prepared to aid data analysis.

sgRNAs were designed to target conserved functional domains that display low computationally predicted off-target scores using the GUIDES tool (http://guides.sanjanalab.org/), preferentially picking sgRNAs that have an A/T nucleotide at the 17^th^ position of the sgRNA sequence. These sgRNA oligo pools were cloned into the LRG expression vector following which the plasmid pool was amplified, purified, and packaged into lentiviruses. To ensure that MTB/TAN-Cas9 cells receive only one sgRNA/cell, the lentiviral library was titered using the GFP selectable marker by flow cytometry (Attune NxT Thermo Fisher) and transduced at an MOI=0.3. Finally, the transduced cells were sorted (MoFlo Astrios, Beckman Coulter Life Sciences) and expanded prior to transplantation in vivo.

All samples harvested from mouse primary tumors and residual lesions were microdissected under a stereoscope and homogenized for genomic DNA extraction using the Quick-DNA Midiprep Plus kit (Zymo Research, Cat. # D4075). sgRNA inserts were amplified by PCR using the Phusion Flash High Fidelity PCR Master Mix (Thermo Fisher, Cat. # F548), followed by barcode and adapter addition. The libraries were then pooled, mixed with 5% PhiX (Thermo Fisher, Cat. # FC-110-3001) and massively parallel sequenced using the MiSeq reagent kit v3 150 cycle (Illumina, Cat. # MS-102-3001) on the MiSeq Instrument (Illumina).

Fastq files containing sequencing reads were demultiplexed using an in-house R script, followed by sgRNA counting using a modified version of a published Python script. Percent abundance was calculated for each sgRNA relative to total read count per sample. Read counts from technical replicates were combined. Diversity in read distribution across sgRNAs in a sample was visualized by plotting the cumulative percentage of reads against descending ranks of sgRNAs by read counts for comparison across different samples and time points. Potential clonal or emerging clonal genes were identified by looking for sgRNAs satisfying at least one of the following: 1) percent abundance in a sample is >5%, abundance is more than 1.5x interquartile range above the upper quartile, or 3) abundance in a sample is significantly above the levels in primary tumor samples as determined using edgeR with a fold-change cutoff of 2 and an FDR cutoff of 0.1.

## Histology

For in vitro immunofluorescence/labeling studies, MTB/TAN-derived primary tumor cells were cultured on glass coverslips (Bellco Glass Inc., Cat. # 1943–010015A) and treated with 5μM EdU for 2h before harvest. Coverslips containing cells were fixed in 4% paraformaldehyde for 15 min at room temperature. Cells were then permeabilized in PBS containing 0.5% Triton X-100 for 20 min at room temperature followed by wash steps in 3% bovine serum albumin in 1x PBS before proceeding with the immunofluorescence protocol.

For in vivo immunofluorescence/labeling studies, inguinal #4 mammary fat pads containing primary tumors, residual lesions, or recurrent tumors were dissected and fixed overnight in 4% paraformaldehyde overnight at 4°C. Samples were thoroughly washed and dehydrated prior to paraffin embedding and sectioning at 5μm. Slides were prepared by serial deparaffinization and rehydration followed by PBS washes and antigen retrieval in R-Buffer A (Electron Microscopy Sciences, Cat. # 62706–10).

For labeling studies using coverslips for EdU (Thermo Fisher Scientific, Cat. #C10640) or on tissues for EdU (Sigma Aldrich, Cat. # BCK647-IV-IM-S), samples were processed as per manufacturer’s instructions prior to immunofluorescence.

Immunofluorescence samples were blocked with 5% goat serum in 1x PBS, washed 3 times in 1x PBS and incubated overnight at 4°C with primary antibodies or matched isotype controls diluted in 5% goat serum in 1x PBS (Vector Laboratories, Cat. # BMK-2202) as follows: <u>Primary antibodies</u> - Rat anti-Ki67 (Thermo Fisher Scientific, Cat. # 14–5698; 1:100), Chicken anti-GFP (Abcam, Cat. # AB13970, 1:600); <u>Isotype controls</u> - Rat IgG (Thermo Fisher Scientific, Cat. # 02–9602), Normal Chicken IgY (Abcam, Cat. # AB-101-C).

After performing 3 washes with 1x PBS, samples were incubated with secondary antibodies at 37°C for 1h as follows: <u>Secondary antibodies</u> - Goat anti-rat IgG Alexa568 (Thermo Fisher Scientific, Cat. # A11077; 1:1000), Goat anti-chicken IgY Alexa488 (Thermo Fisher Scientific, Cat. # A11039, 1:1000).

Following incubation with secondary antibodies, samples were washed 3 times with 1x PBS and incubated with 0.5μg/ml Hoechst 33258 for 10 min at room temperature for nuclear counter staining. Samples were mounted using ProLong gold (Thermo Fisher Scientific, Cat. #P36394) and imaged using a DM 5000B Automated Upright Microscope with a DFC350 FX monochrome digital camera (Leica Microsystems). Images were quantified using QuPath-0.4.4 open source software.

For EdU staining in residual lesions, proportions of D28 residual lesions that were subclinically recurring were compared using Fisher’s exact test.

## Droplet digital PCR

Microdissected lesions from the in vivo experiments or cells harvested from the in vitro dormancy experiment were processed for genomic DNA isolation using the Quick-DNA Midiprep Plus kit (Zymo Research, Cat. # D4075) kit, or the QIAamp DNA mini kit (Thermo Fisher, Cat. # 51304), respectively. To quantify the numbers of GFP^+^ and mCherry^+^ cells, 50ng of genomic DNA/sample was added together with the ddPCR Supermix for Probes (Bio-Rad, Cat. # 1863025) and the probes of interest: GFP (Bio-Rad, Cat. # dCNS372378948), mCherry (Bio-Rad, Cat. # dCNS507694046), and the control ApoB (Bio-Rad, Cat. # dMmuCNS4075944696). Droplets were generated using the AutoDG system (Bio-Rad, Cat. # 1864101) followed by PCR under the following conditions: 95°C for 5 min; 40 cycles at 94°C for 30 sec and 60°C for 1 min; 98°C for 10 min. Droplets were analyzed using the QX200 Droplet Reader (Bio-Rad, Cat. # 1864003). Droplet-derived copy numbers were first normalized to ApoB numbers to normalize input and then converted to cell numbers by using the average copy number derived from singly transduced cells. Differences in GFP% between groups were analyzed by Student’s t-test with Bonferroni correction for multiple comparisons.

## Drug and cellular assays

Erastin (MedChemExpress, Cat. # HY-15763), RSL3 (Selleckchem, Cat. # S8155), and ferrostatin-1 (MedChemExpress, Cat. # HY-100579) were reconstituted in DMSO, and then diluted in DMSO and cell medium (either 10% serum with doxycycline for proliferating cells, 1% serum without doxycycline for dormant cells) prior to administration by changing media on the cells of interest. Following the intended treatment length, for an MTT assay (Promega CellTiter 96 Non-radioactive Cell Proliferation Assay, Cat. # G4000), 15 uL of MTT dye was added per well of a 96-well plate. Following 3hr incubation, the reaction was stopped with 100uL of stop solution per well, incubated at 4C overnight, and then read at 570nm and 630nm absorbance the next day. For microscoping imaging, 5uL NucBlue Live ReadyProbes Reagent (Hoechst 33342) (ThermoFisher Scientific, Cat. # R37605) was added to each 100uL well 1hr prior to imaging on EVOS m7000 microscope. When using, 5uL NucGreen Dead 488 ReadyProbes Reagent (ThermoFisher Scientific, Cat. # R37109) was added to each 100uL well 15-30 min prior to imaging. Statistical significance of differences in apparent tumor cell count were calculated using Student’s t-test with Bonferroni correction for multiple testing.

GSH/GSSG ratio was calculated using the GSH/GSSG assay kit (MedChemExpress, Cat. # HY-K0311) according to manufacturer’s instructions. Proliferating cells were harvested two days after plating and dormant cells were harvested on D14. Briefly, cells were scraped in PBS, resuspended in extraction buffer, sonicated and centrifuged. Supernatant was stored at - 80C for 6 days prior to running the assay. Total and oxidized glutathione were measured, and GSH/GSSG was subsequently calculated.

NADP+/NADPH and NAD+/NADH ratio were calculated using Promega NADP/NADPH-Glo (Cat. # G9081) and NAD/NADH-Glo (Cat. # G9071) according to manufacturer’s instructions for measuring NAD(P)+ or NAD(P)H individually. Standard curves were made using purified NADP+ (Sigma, Cat. # N8035) or NAD+ (Sigma, Cat. # N8285). Ratios were measured in proliferating, D7, and D14 dormant cells.

Liperfluo assay was done according to manufacturer’s protocol in proliferating and dormant D10 cells. Cells were treated with DMSO, 20uM erastin, or 20uM erastin and 20uM ferrostatin for 72hr prior to assay. Cells were washed twice with Hank’s Balanced Salt Solution (ThermoFisher Scientific, Cat. # 14175095) then received 10uM Liperfluo in HBSS. Cells were incubated in Liperfluo for 30 min at 37C, then washed once with HBSS, trypsinized for 5 min with 0.05% trypsin (proliferating cells) or 0.25% trypsin (dormant cells), and then analyzed on Attune NxT flow cytometry (Thermo Fisher). Results were analyzed in FlowJo software.

## Human data analysis

Ferroptosis pathway mutations were analyzed on WES in 130 paired primary and metastatic breast tumors from several different datasets and compared via one-sided Wilcoxon signed-rank test. The distribution of low- and high-level CN gains and losses in *PNPLA2*, *GOT1*, *GPX4* was calculated in the METAMORPH (56) data set’s primary tumors (n = 28) and metastases (n = 66). The p-value indicates one-sided Wilcoxon rank-sum test that compares the CN values between primary and metastatic tumors. OS and RFS for patients with *PNPLA2*, *GOT1*, or *GPX4* copy number change was assessed in the METABRIC cohort (57) by Kaplan- Meier analysis.

---

**Supplementary Figure 1.** : Dotplot results of CRISPR screening display multiple methods of guide analysis Dotplots showing the relative abundance (left), interquartile range (IQR) distance (left-center), false discovery rate (right-center), and log2 fold change (right) for each of the genes of interest in the CRISPR screen. Individual guides are identified by color and timepoints from the screen are organized vertically. a. *PNPLA2*, b. *GOT1*, c. *GPX4*.

**Supplementary Figure 2:** ***ICE analysis confirms guides used target genes of interest*** Inference of CRISPR edits (ICE) analysis for: a. *PNPLA2g4* b. *PNPLA2g5* c. *GOT1g1* d. *GOT1g5* e. *GPX4g5* f. *GPX4g8*. Output shows percentage of indels and predicted knockout efficiency (“Knockout-Score”). Wild-type sequence is denoted by an orange + sign. Vertical dotted line denotes the Cas9 cut site.

**Supplementary Figure 3:** Recurrence-free survival from in vivo competition experiment provides evidence that ferroptosis regulators accelerate recurrence Kaplan-Meier curves for: a. *PNPLA2g4* and *PNPLA2g5*, b. *GOT1g1* and *GOT1g5*, c. *GPX4g5* and *GPX4g8* of recurrence-free survival compared to sg*Rosa* control in the in vivo competition experiment shown in Fig 4. For each guide listed, that guide comprised ∼20% of the injected pool of cells and was tagged with GFP and was in competition with a Rosa guide tagged with mCherry comprising the remainder of the cell population.

**Supplementary Figure 4:** Longer term erastin treatment in dormant and proliferative cell states mirrors 24hr treatment Results of MTT assay after treating: a,b. proliferative iHer2 cells, c,d. iHer2 cells at D7 of dormancy, e,f. iHer2 cells at D14 of dormancy with increasing doses of erastin for 48hr (a,c,e) or 72hr (b,d,f). MTT absorbance in each experiment, as a proxy for cell viability, is normalized to its own vehicle control (equal volume DMSO).

**Supplementary Figure 5:** Dormant residual tumor cell number increases in response to high-dose erastin a. MTT assay comparing proliferative (solid bars) to D7 iHer2 cells (striped bars) with 24 hr of erastin treatment. b. Total cell count as assessed by number of NucBlue positive nuclei in microscopic images taken by EVOS m7000 microscope following 24 hr of erastin treatment for proliferative (solid bars) or D7 iHer2 cells (striped bars). Each cell state is normalized to its respective vehicle control.

**Supplementary Figure 6:** Proliferative iHer2 cell death is partially reversible by ferrostatin-1 a. MTT results of proliferative iHer2 cells treated with erastin, RSL3, ferrostatin, or a combination of a ferroptosis inducer and ferrostatin-1 for 24hr. b. MTT results of proliferative iHer2 cells treated with multiple doses of erastin and ferrostatin-1 for 24hr. c. MTT results of proliferative iHer2 cells treated with multiple doses of RSL3 and ferrostatin-1 for 24hr. d. Live cell count by microscopic imaging of proliferative iHer2 cells treated with ferrostatin-1 for 24 hr followed by RSL3 for 24hr. e. MTT results of D14 dormant iHer2 cells treated with multiple doses of erastin and ferrostatin-1 for 24hr. f. MTT results of D14 dormant iHer2 cells treated with multiple doses of RSL3 and ferrostatin-1 for 24hr.

**Supplementary Figure 7:** Human breast cancer overall survival associations with PNPLA2, GOT1, and GPX4 parallel recurrence-free survival mouse experiments a-c. Overall survival outcomes for METABRIC patients harboring loss of *PNPLA2* (a), *GOT1* (b), and *GPX4* (c) across all tumors. d-f. Overall survival outcomes for METABRIC patients harboring loss of *PNPLA2* (d), *GOT1* (e), and *GPX4* (f) in only HR+ tumors.

**Supplementary Figure 8:** PNPLA2 and GPX4 expression in human breast cancer patients is associated with ER+HER2- disease and lower tumor grade. a. Relative expression of *PNPLA2* stratified by ER/HER2 status. b. Relative expression of *PNPLA2* stratified by tumor grade. c. Relative expression of *GOT1* stratified by ER/HER2 status. d. Relative expression of *GOT1* stratified by tumor grade. e. Relative expression of *GPX4* stratified by ER/HER2 status. f. Relative expression of *GPX4* stratified by tumor grade.

**Supplementary Figure 9:** Human breast cancer relapse data supports role of sgPNPLA2, sgGOT1, and sgGPX4 in accelerating recurrence a-c. Human breast cancer patient gene expression associations with 5-year relapse for *PNPLA2* (a), *GOT1* (b), and *GPX4* (c). Green dot indicates association between higher gene expression and better RFS; red dot indicates association between higher gene expression and worse RFS. Increased number of dots indicates increased statistical significance.

