## Supplementary Figures for "Loss of Ferroptosis-suppressors Accelerates Breast Cancer Recurrence by Promoting Dormancy Exit"

### Supplementary Figure 1

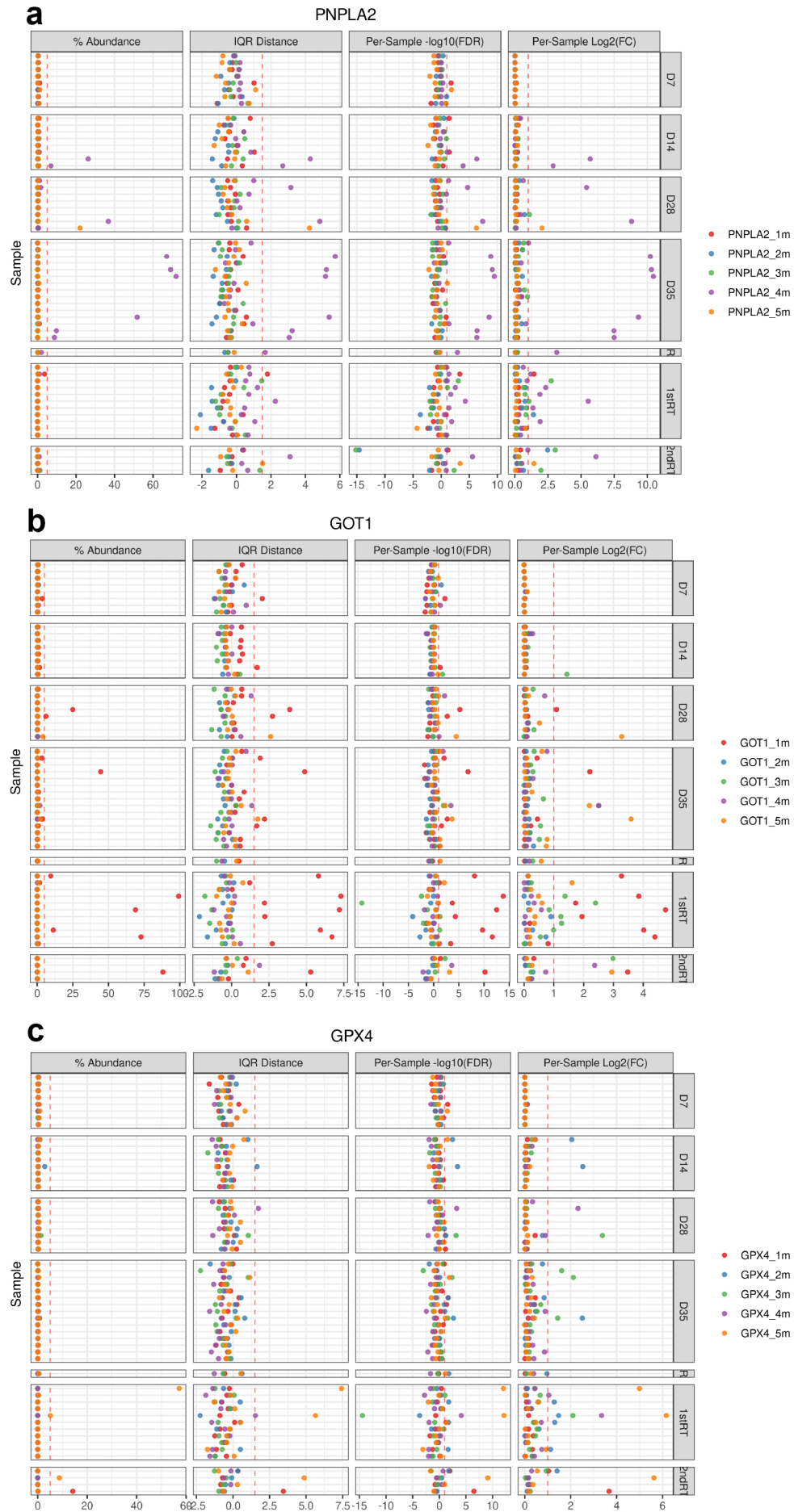

### Supplementary Figure 2

a

#### PNPLA2g4

| Status | Guide Target | PAM Sequence | Indel % | Model Fit (R<sup>2</sup>) | Knockout-Score | Nuclease |
| --- | --- | --- | --- | --- | --- | --- |
| ✓ Succeeded | TCCGCTCTGAACACGAGTCA | GGG | 87 | 0.91 | 87 | spcas9 |

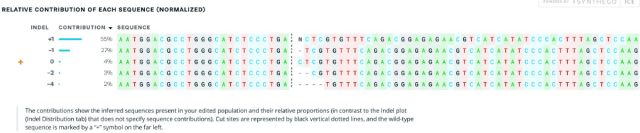

b

#### PNPLA2g5

| Status | Guide Target | PAM Sequence | Indel % | Model Fit (R<sup>2</sup>) | Knockout-Score | Nuclease |
| --- | --- | --- | --- | --- | --- | --- |
| ✓ Succeeded | AGAGGCGGTAGAGATTGCGA | AGG | 84 | 0.95 | 76 | spcas9 |

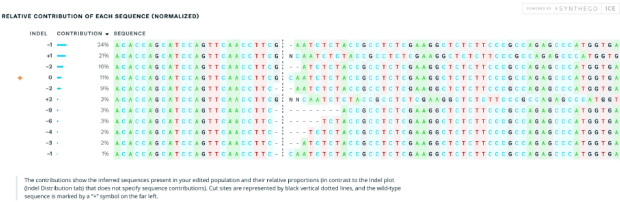

c

#### GOT1g1

| Status | Guide Target | PAM Sequence | Indel % | Model Fit (R<sup>2</sup>) | Knockout-Score | Nuclease |
| --- | --- | --- | --- | --- | --- | --- |
| ✓ Succeeded | AGCTTAAAGACCAAGACCG | AGG | 61 | 0.92 | 54 | spcas9 |

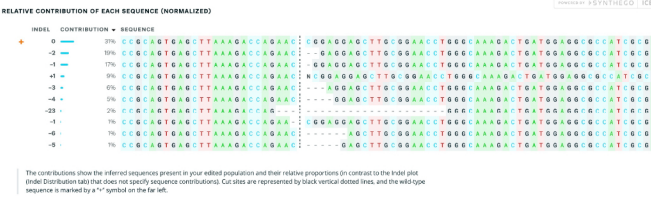

d

#### GOT1g5

| Status | Guide Target | PAM Sequence | Indel % | Model Fit (R<sup>2</sup>) | Knockout-Score | Nuclease |
| --- | --- | --- | --- | --- | --- | --- |
| ✓ Succeeded | GATGTGAGACCAAGTCCCGG | GGG | 66 | 0.95 | 64 | spcas9 |

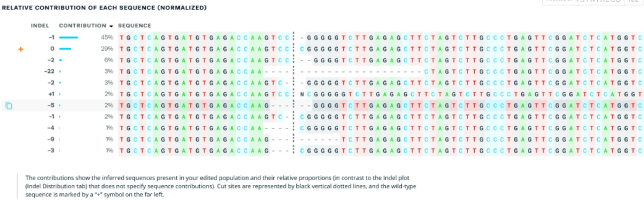

e

#### GPX4g5

| Status | Guide Target | PAM Sequence | Indel % | Model Fit (R<sup>2</sup>) | Knockout-Score | Nuclease |
| --- | --- | --- | --- | --- | --- | --- |
| ✓ Succeeded | CATCAAAATGGAACCTTTACCA | AGG | 80 | 0.97 | 67 | spcas9 |

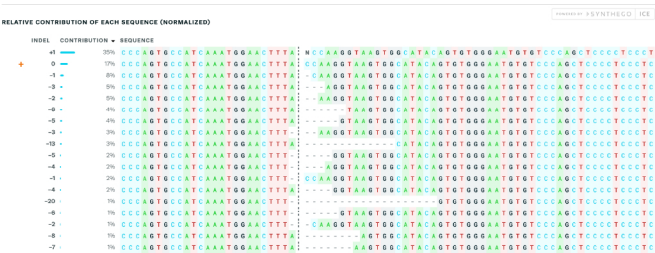

f

#### GPX4g8

| Status | Guide Target | PAM Sequence | Indel % | Model Fit (R<sup>2</sup>) | Knockout-Score | Nuclease |
| --- | --- | --- | --- | --- | --- | --- |
| ✓ Succeeded | ACCAACGTGGCTCGCAATG | AGG | 87 | 0.95 | 87 | spcas9 |

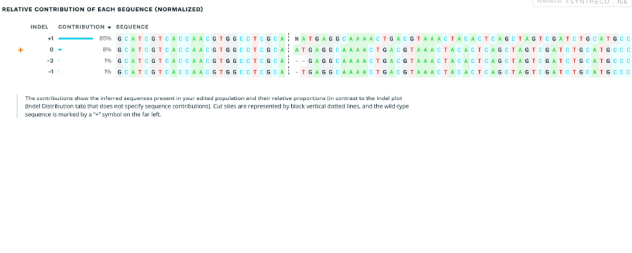

#### Supplementary Figure 3

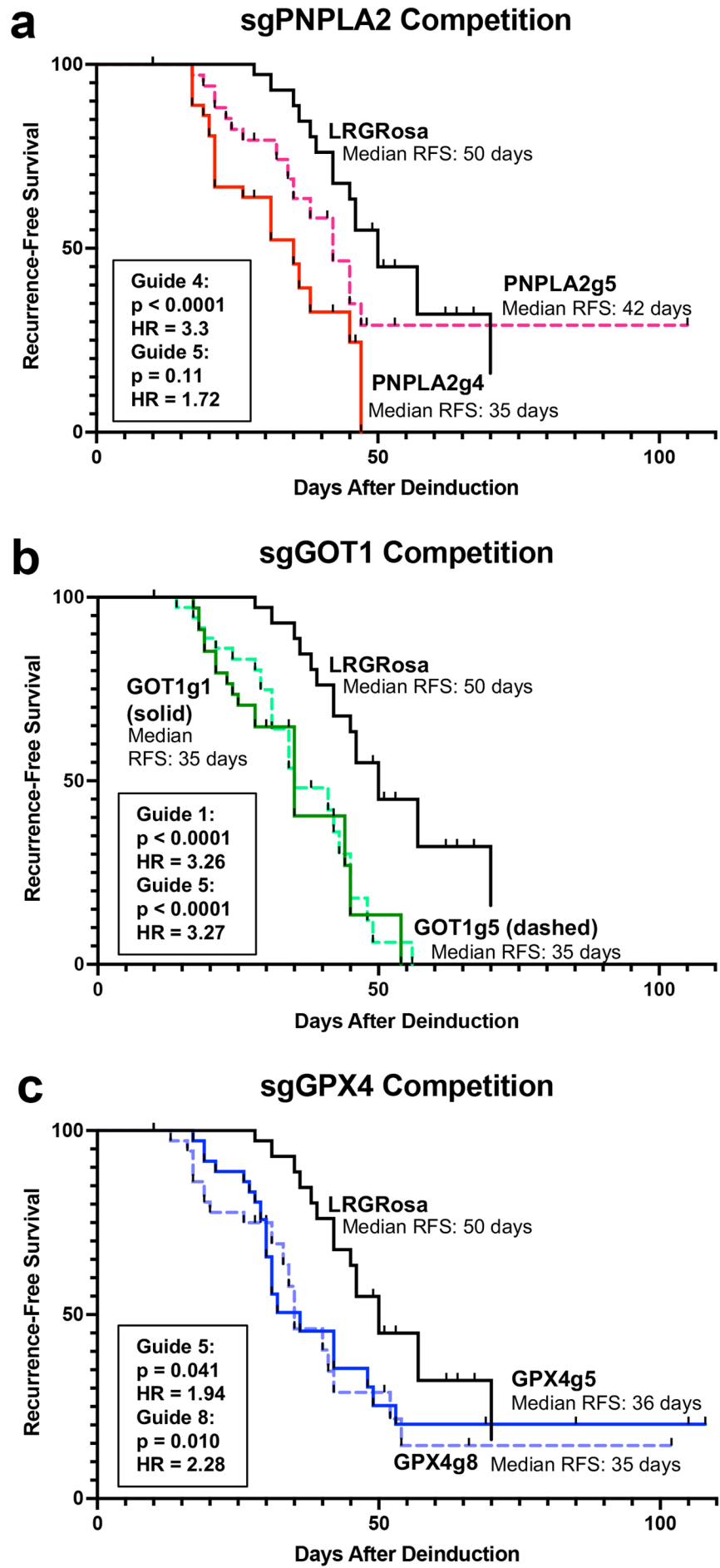

#### Supplementary Figure 4

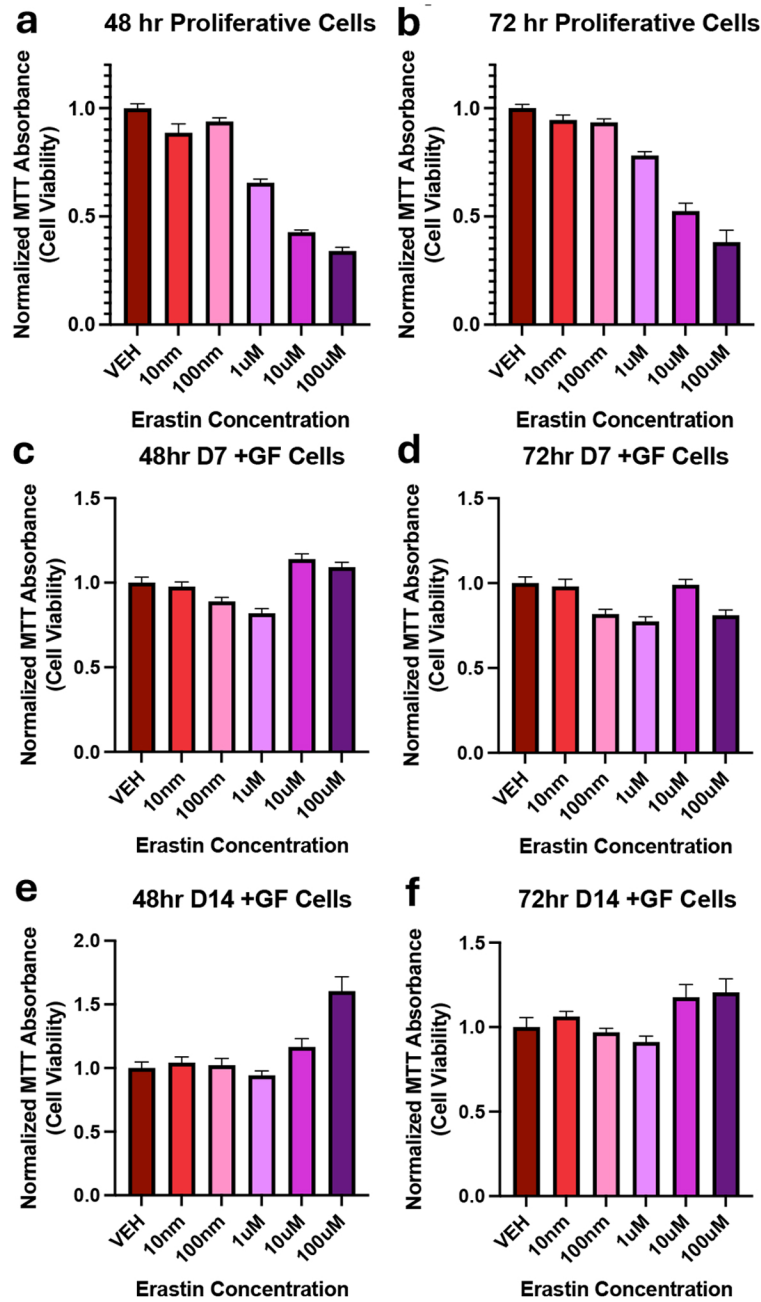

#### Supplementary Figure 5

**a** MTT Assay Proliferative vs. Dormant

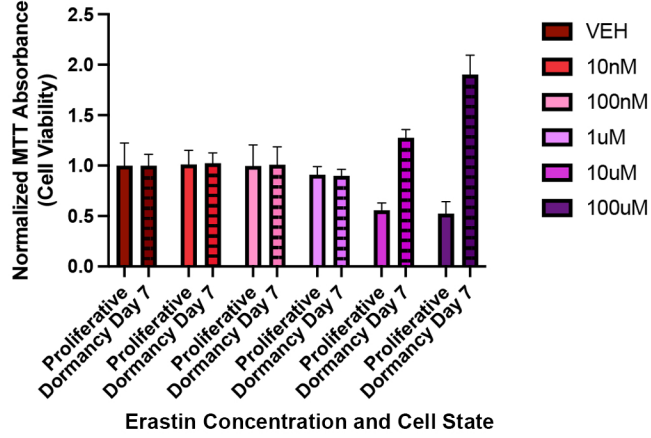

**b** NucBlue Cell Count

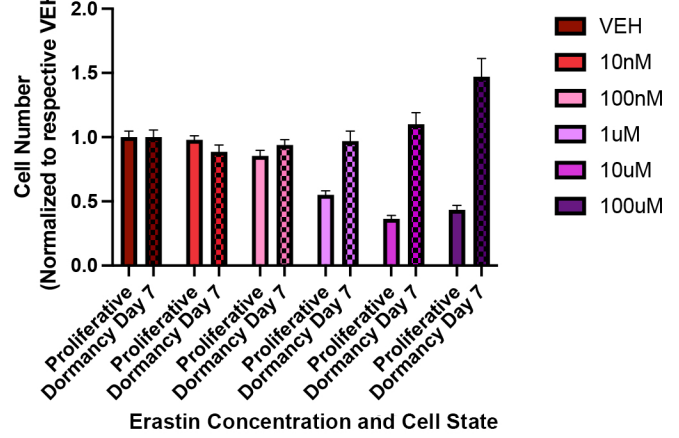

### Supplementary Figure 6

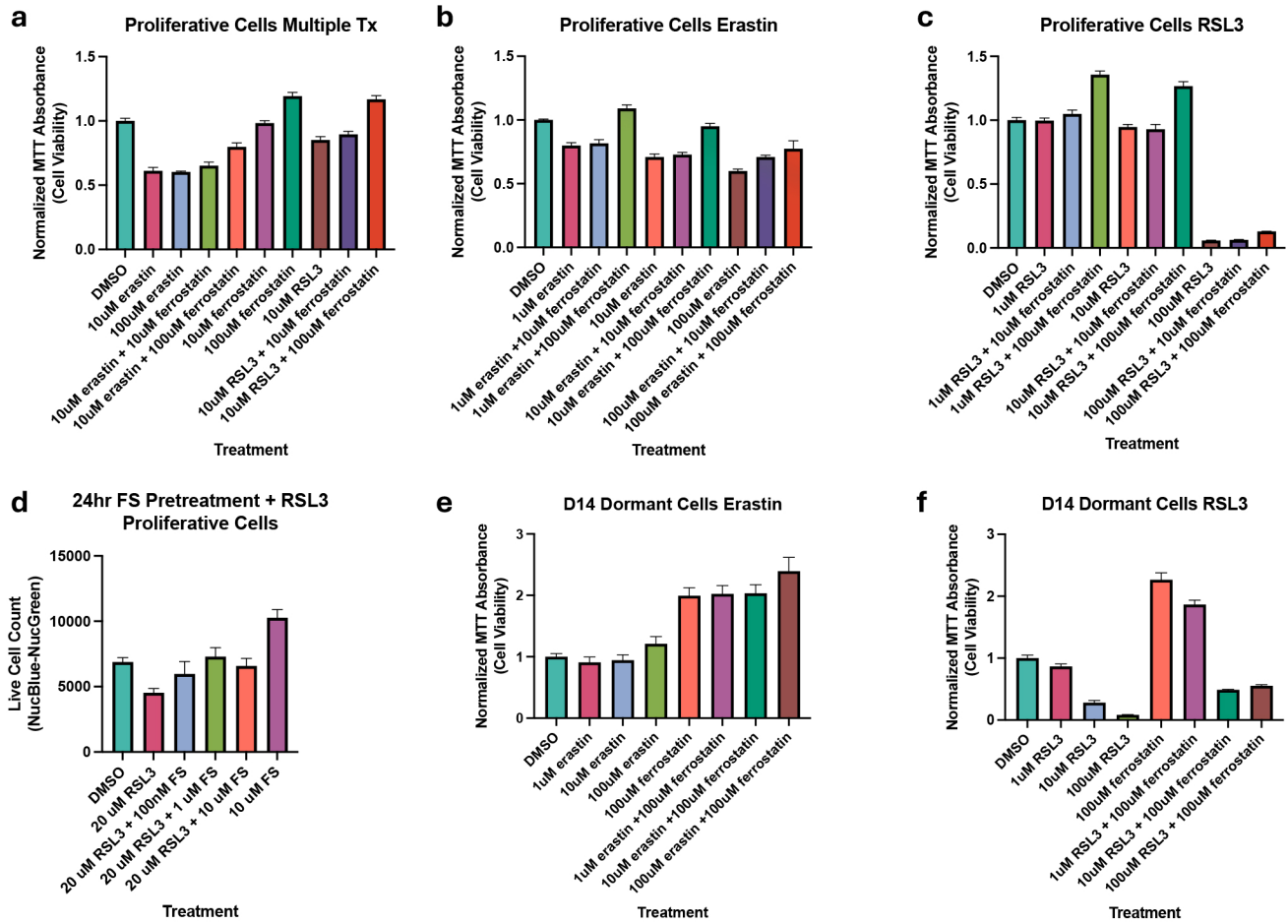

#### Supplementary Figure 7

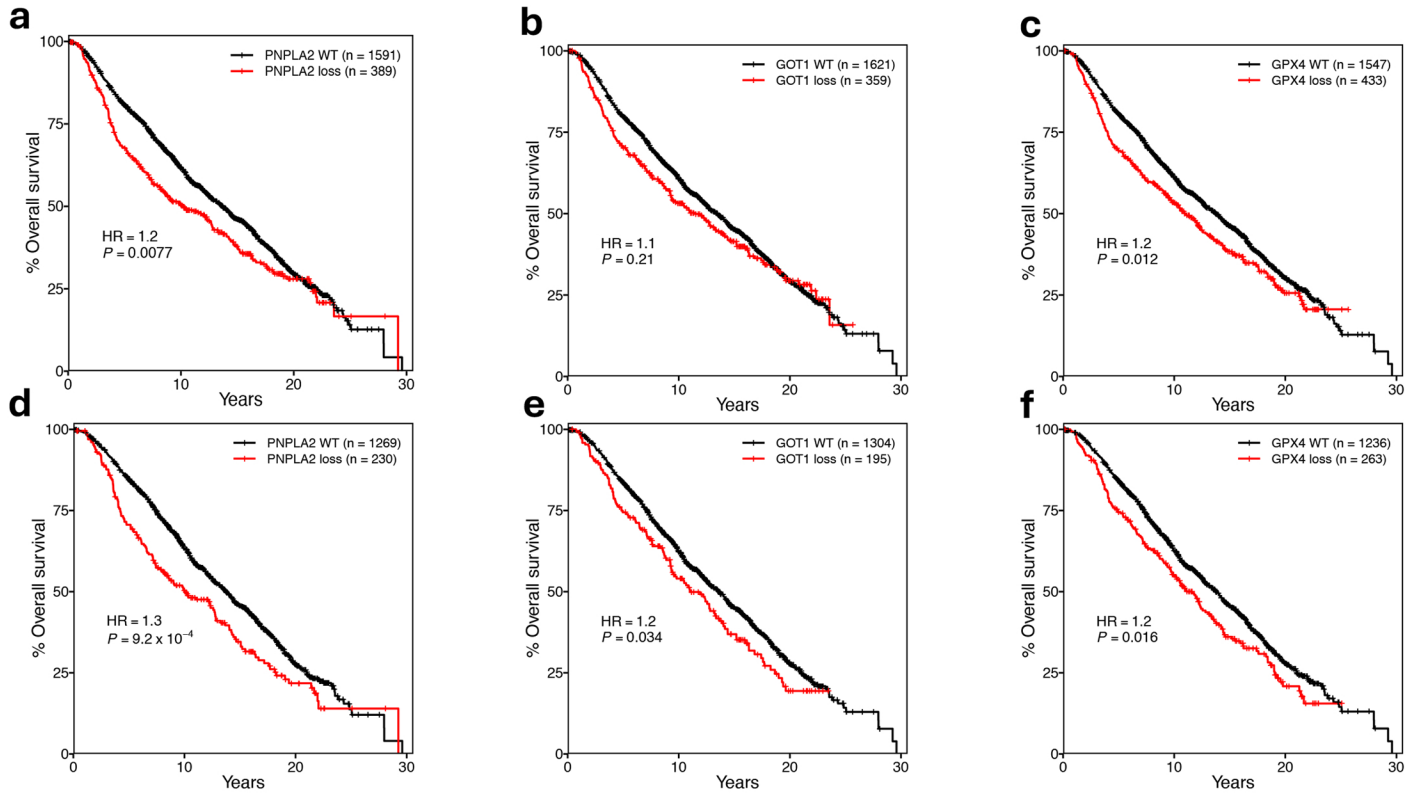

#### Supplementary Figure 8

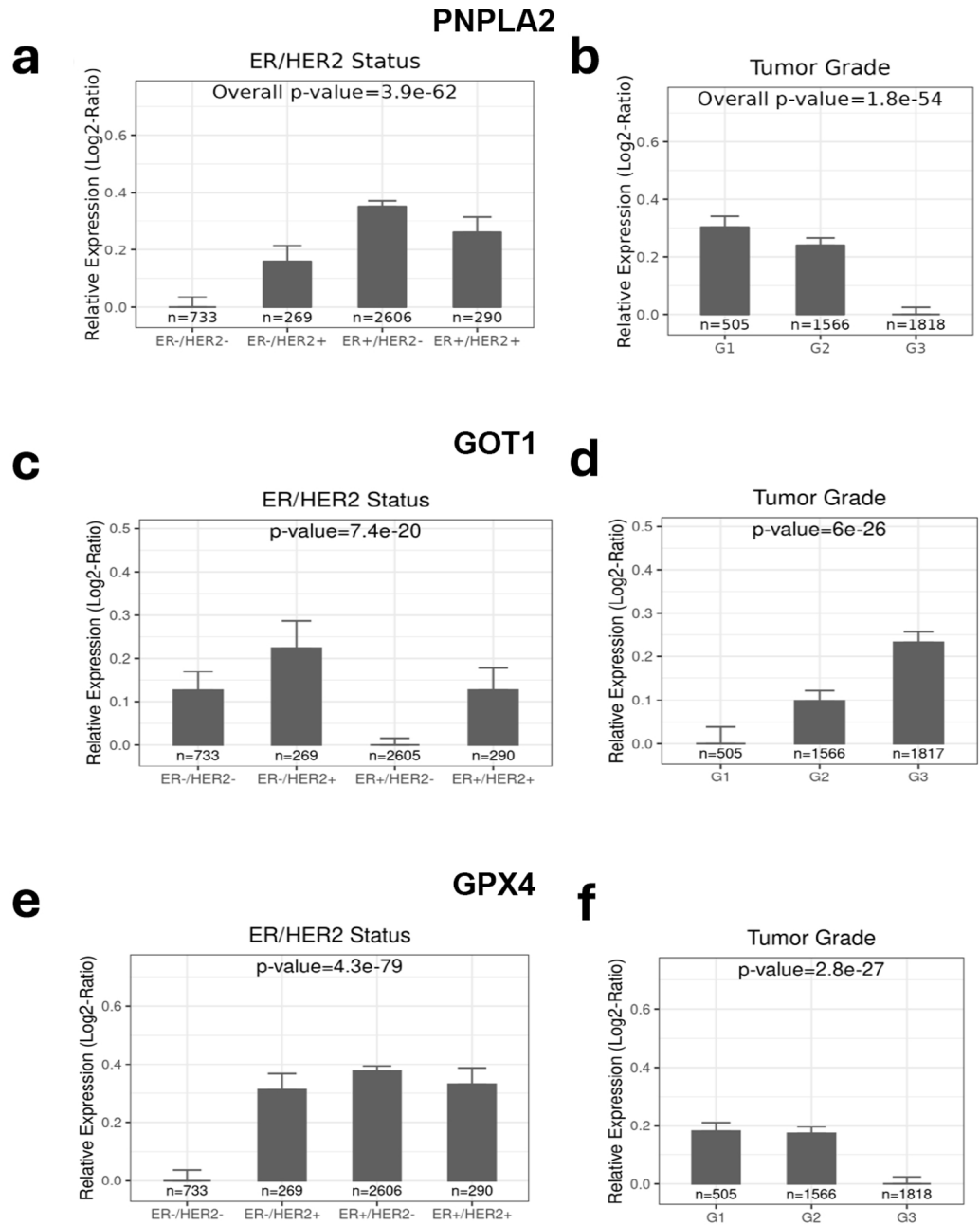

### Supplementary Figure 9

**a** **PNPLA2**
Human Breast Cancer 5-Year Relapse-Free Survival

| Strata | Mixed Tx |  |  | No Systemic Tx |  |  |
| --- | --- | --- | --- | --- | --- | --- |
|  | Cox-Cont | C-Index | Cox-Bin | Cox-Cont | C-Index | Cox-Bin |
| All | • | ••• | ••• | ••• | ••• | ••• |
| ER+ | • | •• | •• | •• | •• |  |
| ER- |  |  |  |  |  |  |
| HER2+ |  |  |  |  |  |  |
| HER2- | ••• | ••• | ••• | ••• | ••• | ••• |
| ER+/HER2+ |  |  |  |  |  |  |
| ER+/HER2- | ••• | ••• | •• | •• | •• |  |
| ER-/HER2+ |  |  |  |  |  |  |
| ER-/HER2- |  |  |  |  | • |  |
| Node+ |  | • | • |  |  |  |
| Node- | ••• | ••• | ••• | ••• | ••• | ••• |
| G1&G2 |  |  |  |  |  |  |
| G3 |  |  | • |  | • |  |
| Basal |  |  |  |  | • |  |
| ERBB2+ |  |  |  |  |  |  |
| Luminal A |  |  |  | • |  |  |
| Luminal B |  |  |  |  |  |  |
| Normal-like |  |  |  |  |  |  |

Data version: 2012-10-03 04:09:55

**b** **GOT1**
Human Breast Cancer 5-Year Relapse-Free Survival

| Strata | Mixed Tx |  |  | No Systemic Tx |  |  |
| --- | --- | --- | --- | --- | --- | --- |
|  | Cox-Cont | C-Index | Cox-Bin | Cox-Cont | C-Index | Cox-Bin |
| All | ••• | ••• | ••• | • | • |  |
| ER+ | ••• | ••• | •• | • |  |  |
| ER- |  | • |  |  |  |  |
| HER2+ |  |  |  |  |  |  |
| HER2- | ••• | ••• | ••• |  |  |  |
| ER+/HER2+ |  |  |  |  |  |  |
| ER+/HER2- | •• | •• | • |  |  |  |
| ER-/HER2+ |  |  |  |  |  |  |
| ER-/HER2- |  | • |  |  |  |  |
| Node+ | ••• | ••• | ••• |  |  |  |
| Node- | • | • |  | • |  |  |
| G1&G2 | ••• | ••• | ••• | •• | •• | • |
| G3 |  |  |  |  |  |  |
| Basal | • | •• |  |  |  |  |
| ERBB2+ |  |  |  |  |  |  |
| Luminal A |  |  |  |  |  |  |
| Luminal B |  |  |  |  |  |  |
| Normal-like |  |  |  |  |  |  |

Data version: 2012-10-03 04:09:55

**c** **GPX4**
Human Breast Cancer 5-Year Relapse-Free Survival

| Strata | Mixed Tx |  |  | No Systemic Tx |  |  |
| --- | --- | --- | --- | --- | --- | --- |
|  | Cox-Cont | C-Index | Cox-Bin | Cox-Cont | C-Index | Cox-Bin |
| All | • | ••• | •• | •• | ••• | ••• |
| ER+ |  |  |  | • | • | • |
| ER- |  |  |  |  |  |  |
| HER2+ |  |  | • |  |  |  |
| HER2- | ••• | ••• | •• | •• | ••• | •• |
| ER+/HER2+ |  |  |  |  |  |  |
| ER+/HER2- |  |  |  |  |  |  |
| ER-/HER2+ |  |  |  |  |  |  |
| ER-/HER2- |  |  |  |  |  |  |
| Node+ |  |  |  |  |  |  |
| Node- | •• | ••• | •• | •• | ••• | ••• |
| G1&G2 |  |  |  |  |  |  |
| G3 | • | •• | • | • | • |  |
| Basal |  |  | • |  |  |  |
| ERBB2+ |  |  |  |  |  |  |
| Luminal A |  |  |  |  |  |  |
| Luminal B | •• |  |  | • |  |  |
| Normal-like |  |  |  |  |  |  |

Data version: 2012-10-03 04:09:55
